# The coordinated regulatory roles of two LysR-Type Transcriptional Regulators balance chorismate and protocatechuate partition in *Listeria* organisms

**DOI:** 10.1101/2025.04.08.647683

**Authors:** Kevin Xue, Cindy Yookyung Hong, Elina Kadriu, Dinesh Christendat

## Abstract

*Listeria monocytogenes* is an economically deleterious foodborne pathogens that continually challenges the global food supply chain. *Listeria* species in general, synthesize protocatechuate from saprophytically-derived quinate and shikimate utilizing a novel class of bacterial dehydroshikimate dehydratase. Paradoxically, *Listeria* species are unable to metabolically utilize protocatechuate, as such, it was proposed that this compound is used as a currency to *Listeria* interactions with other microorganisms to improve their environmental proliferation. Therefore, an understanding of the regulatory mechanism for the metabolic pathway for protocatechuate biosynthesis is of great importance. Two LysR Type Transcriptional Regulators (LTTR), annotated QuiR and in this study QuiR2, are found upstream of genomic operons, *qui1* and *qui2,* which transcribe genes for protocatechuate synthesis. QuiR, has been shown to activate the expression of genes from both operons with shikimate as a coinducer. However, the role of QuiR2, Lmo2233, is not clear. In this study, we conducted structural, biochemical and bioinformatics analyses of QuiR2 and demonstrated that it functions as a negative regulator of protocatechuate biosynthesis in *Listeria* species. Moreover, we determined that protocatechuate functions in modulating QuiR2 repressive properties through our mobility shift assay and LacZ reporter activity studies. Furthermore, phylogenetic analyses reveal that QuiR2 clusters closely but independently from QuiR thus supporting their distinct regulatory roles. We propose that QuiR2 prevents metabolic commitment of dehydroshikimate to protocatechuate when elevated and in limiting shikimate condition. In this study we revisited the biological role of the shikimate pathway in microbes and demonstrated that in addition to it producing chorismite for aromatic compound metabolism it is also important in allowing organisms to shuttle shikimate and quinate to produce protocatechuate which can be used as an energy source and more importantly in *Listeria* it is used to facilitate microbial interactions.

## Introduction

LysR-type transcriptional regulators (LTTRs) belong to a large family of ubiquitous bacterial proteins which regulate the expression of genes with diverse biological functions. These includes genes for proteins involved in various cellular processes, including metabolism, cell division, quorum sensing, virulence, cell adhesion, and secretion (Maddocks and Oyston, 2008; Monferrer *et al*., 2010). The three-dimensional structure of these transcriptional regulators is characterized by a prototypical N-terminal winged helix-turn-helix DNA binding domain (DBD) and a C-terminal effector binding domain (EBD). LTTR conformations rely on their oligomerization states where dimeric LTTRs interact with one another to form a homo-tetramer to facilitate DNA binding (Giannopoulou *et al*., 2021). Additionally, effector binding influences the tetrameric conformation of LTTRs, thus altering their ability to regulate gene expression by bending the DNA and changing accessibility of the RNA polymerase.

Similarly to Isocitrate Lyase-Type Regulators (IclRs), LTTRs regulate gene expression for proteins that are involved in the metabolism of aromatic compounds including environmental pollutants (Tropel and Van Der Meer, 2004). Thus, LTTRs provide an attractive avenue for engineering organisms to utilize aromatic compounds. In addition, LTTRs regulate gene expression for complete metabolic pathways that are involved in the synthesis of signaling molecules and other biologically relevant compounds. For example, in *Listeria monocytogenes*, the QuiR LTTR induces gene expression for enzymes involved in the protocatechuate biosynthesis pathway (Prezioso et al., 2018). Protocatechuate biosynthesis in microorganisms occurs by diverting intermediates from the shikimate pathway (Xue et al., 2020). Additionally, protocatechuate is a common intermediate in the microbial degradation of aromatic compounds, such as lignins, phthalates, and polycyclic aromatic hydrocarbons from natural and anthropogenic sources (Harwood and Parales, 1996; Ni *et al*., 2013). In microorganisms, protocatechuate precedes the β-ketoadipate pathway, which converts protocatechuate into succinate for energy through the tricarboxylic acid cycle. This mode of protocatechuate utilization in *Pseudomonas putida* allows it to utilize either quinate or shikimate as a sole carbon energy source (Peek *et al*., 2017). This metabolic route provides *P. putida* a strategy to utilize environmentally derived compounds for energy. Alternatively, protocatechuate serves as a precursor for the biosynthesis of siderophores in a subset of anaerobic soilborne and oceanic microorganisms (Barbeau *et al*., 2002; Fox *et al*., 2008; Pfleger *et al*., 2008). Enigmatically, protocatechuate utilization in *Listeri*a is not apparent as production of this benzoic acid derivative has not been shown to lead directly to energy production (Simon *et al*., 1995; Prezioso *et al*., 2018). These findings were supported by bioinformatic analyses, which showed that canonical genes for protocatechuate utilization pathways are absent in *Listeria* genomes (Prezioso et al., 2018). Further, shikimate-treated cultures accumulated protocatechuate without showing signs of protocatechuate utilization. Rather, these cultures have an accelerated death phase, suggesting a toxic buildup of protocatechuate derivatives (Prezioso et al., 2018; Xue et al., 2020).

In *Listeria,* QuiR is known to activate protocatechuate biosynthesis by inducing expression of genes from the *qui1* and *qui2* operons in a shikimate-dependent manner (**Figure 1)**. Quinate dehydrogenase (QDH), YdiB, dehydroquinate dehydratase (DHQD), AroD, and shikimate dehydrogenase (SDH) together with SdhD, shuttle shikimate pathway intermediates toward protocatechuate synthesis. Additionally, it was shown that the SdhD enzyme converts shikimate into dehydroshikimate; the precursor for protocatechuate. The final biosynthetic step involves the conversion of dehydroshikimate to protocatechuate by a dehydroshikimate dehydratase (DSD) enzyme, QuiC2 (Prezioso et al., 2018; Xue et al., 2020).

**Figure 1:**
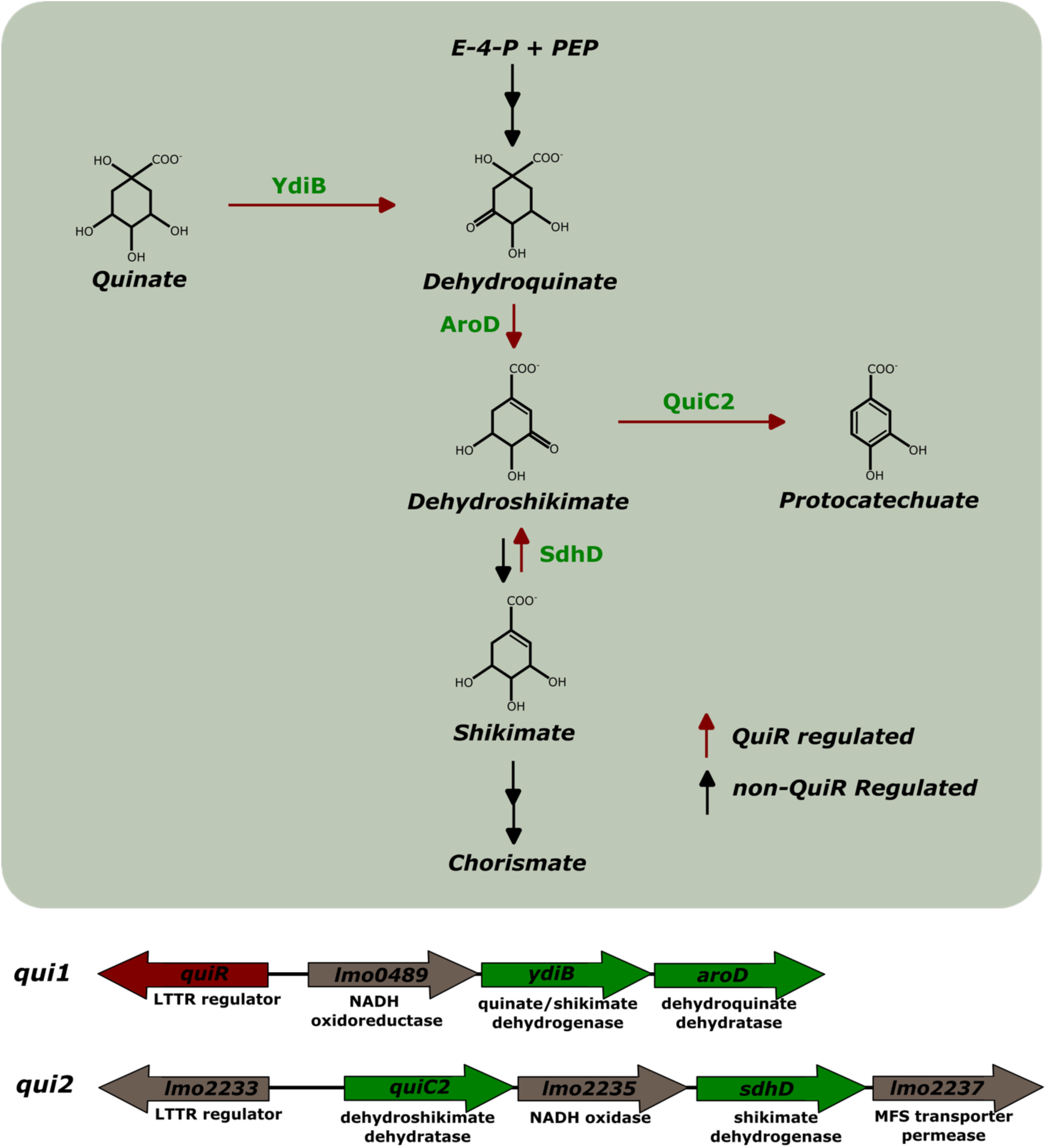
Schematic of the protocatechuate biosynthesis pathway and its regulatory operons. The protocatechuate biosynthesis pathway (top) is facilitated by enzymes encoded by *qui1* and *qui2* operons. Enzymatic reactions catalyzed by these operons are shown by red arrows. Intermediary reactions involving multiple steps are marked by double arrows. LysR type transcription regulators (LTTRs), *quiR* and *lmo2233*, are found upstream of the *qui1* and *qui2* operons (bottom). Each operon consists of previously characterized (green) and uncharacterized (brown) genes.

It has been established that QuiR positively regulates the expression of genes including QuiC2 to drive protocatechuate biosynthesis (Prezioso et al., 2018; Xue et al., 2020). Metabolite commitments to this pathway has been hypothesized to mediate microbial interactions. However, a detail regulatory model has been elusive and requires further investigation. Bioinformatics analysis identified a second LTTR, Lmo2233, encoded divergently upstream of the *qui2* operon. This LTTR gene is positioned immediately upstream of the *qui2* operon indicating that it may also participate in regulating gene expression for this operon (Maddocks and Oyston, 2008; Monferrer *et al*., 2010).

In this study, we examined the regulatory role of Lmo2233 (which we annotate as QuiR2), to further explore the role of protocatechuate production in *Listeria*. Through the use of bioinformatic, biochemical, and microbiological approaches, we established that QuiR2 is a negative regulator of the *qui1* and *qui2* operons’ gene expression. Docking analyses and electrophoretic mobility shift assays (EMSA) indicate that protocatechuate interacts with QuiR2 and modulate gene expression for both *qui1* and *qui2* operons.

## Results

### Phylogenetic analyses of QuiR and QuiR2

In order to investigate the relationship between QuiR2 with characterized LTTRs, phylogenetic analysis was conducted with groups of functionally distinct LTTRs having high sequence similarity to QuiR2. These groups of LTTR include: AaeR which functions in aromatic carboxylate efflux, CrgA which regulates capsule biosynthesis, CatM which controls catechol metabolism, BenM which regulates benzoate metabolism, AmpR which facilitates beta-lactamase production, QuiR which activates protocatechuate biosynthesis, and QuiR2 whose function is reported in this study (**Figure 2)** (Deghmane *et al*., 2000; Van Dyk *et al*., 2004; Ezezika *et al*., 2006; Vadlamani *et al*., 2015; Prezioso *et al*., 2018). A maximum likelihood analysis of these LTTRs produces five major clusters which are supported by high bootstrap values, with the lowest being 0.807. Within these major groups, QuiRs and QuiR2s are closely, but independently, clustered. On the other hand, BenM and CatM are intertwined and clustered together. Due to their high sequence similarity, CrgA regulators clustered together, but they are involved in distinct regulatory processes. These LTTRs are related to diverse metabolic pathways such as cysteine biosynthesis, glycine cleavage, pilus and cell wall biosynthesis, and aromatic compound metabolism (Deghmane et al., 2000). The close clustering of QuiRs and QuiR2s may indicate that they function in closely related regulatory pathways.

**Figure 2:**
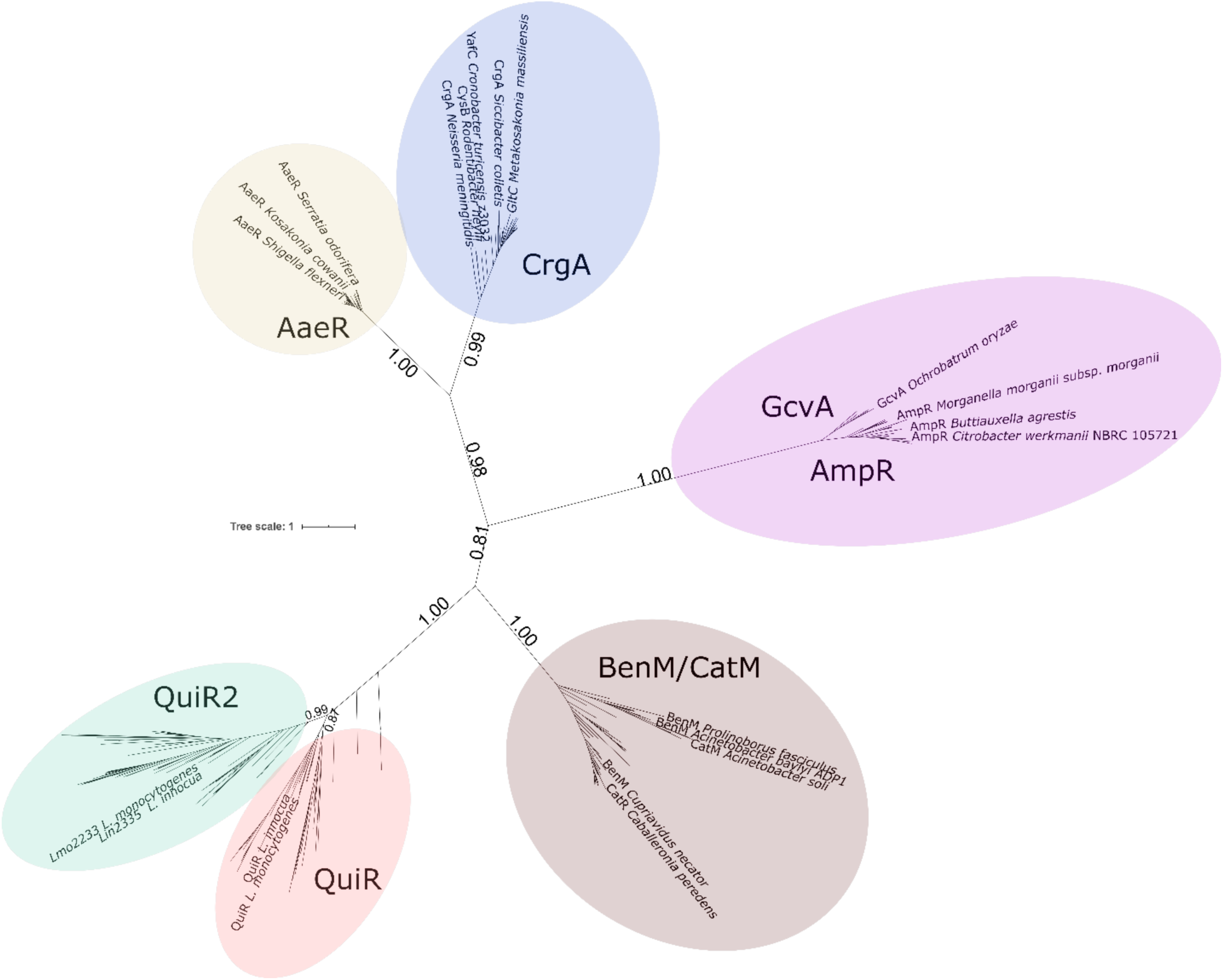
Phylogenetic tree of various LTTRs. Simplified JTT-matrix based maximum likelihood tree of 761 LTTR sequences queried from 7 functional LTTR groups: AaeR, CrgA, AmpR, BenM, CatM, QuiR, and QuiR2. Phylogenetic analysis was performed using the program FastMPTree from XSEDE with 1000 bootstraps. A minimum of 0.81 bootstrap values supports each major branch of the tree. Labeling was performed using ITOL and Inkscape. The full version is included in Figure **S1**.

### Structural analysis of QuiR2 effector binding domain

The structure of the QuiR2 EBD with a bound ligand was attempted by co-crystallizing the protein with a number of ligands that have similar chemical structure to shikimate. The diffraction data from these crystals all yielded the structure of the apoprotein. The protein was crystallized in space group P3_1_2_1_ and the structure was solved to a maximum resolution of 2.52 Å (**Figure** 3**A**). The QuiR2 EBD protein crystal structure was determined by molecular replacement with the use of QuiR crystal structure (Lmo0488; PDB ID: 5TED) as the template. The fact that QuiR consists of two independently folded structural domains prevented initial attempts to obtain a molecular replacement solution. It was thought that the conformational differences between these two domains prevented a successful molecular replacement operation. Therefore, the individual structural domains were separated and used in independent initial molecular replacement searches. The MR solution was associated with high scoring statistics with either domain **(Table 1).** Most of the amino acid residues were fitted and refined during the first round of the model building. The remaining residues were manually placed, and multiple rounds of refinements produced a high-quality model for QuiR2, PDB ID: 9O0T, with R_free_ and R_work_ values of 0.27040 and 0.2196 respectively **(Table 1)**. The overall structural topology of QuiR2 closely resembles QuiR except that the two structural domains are further apart, compared to their organization in QuiR. For example, the distance between QuiR Phe196 and Ile98 is 9.5 Å whereas the corresponding Phe107 and Leu9 are further 14.7 Å apart in QuiR2 **(Figure 3B, C)**. Analysis of the two domains, RD1 (residues 1-75, 177-208) and RD2 (residue 76-176), indicated that they both consist of a *α*/*β* Rossmann fold, with helices surround the central *β*-sheet. Most *β*-strands were oriented parallel to each other. RD1 and RD2 are linked by two antiparallel *β*-strands (residues 156-161, 262-271). Superimpositions of QuiR2 with available LTTR structures were used to identify the primary binding and the likely secondary binding sites of this protein.

**Figure 3:**
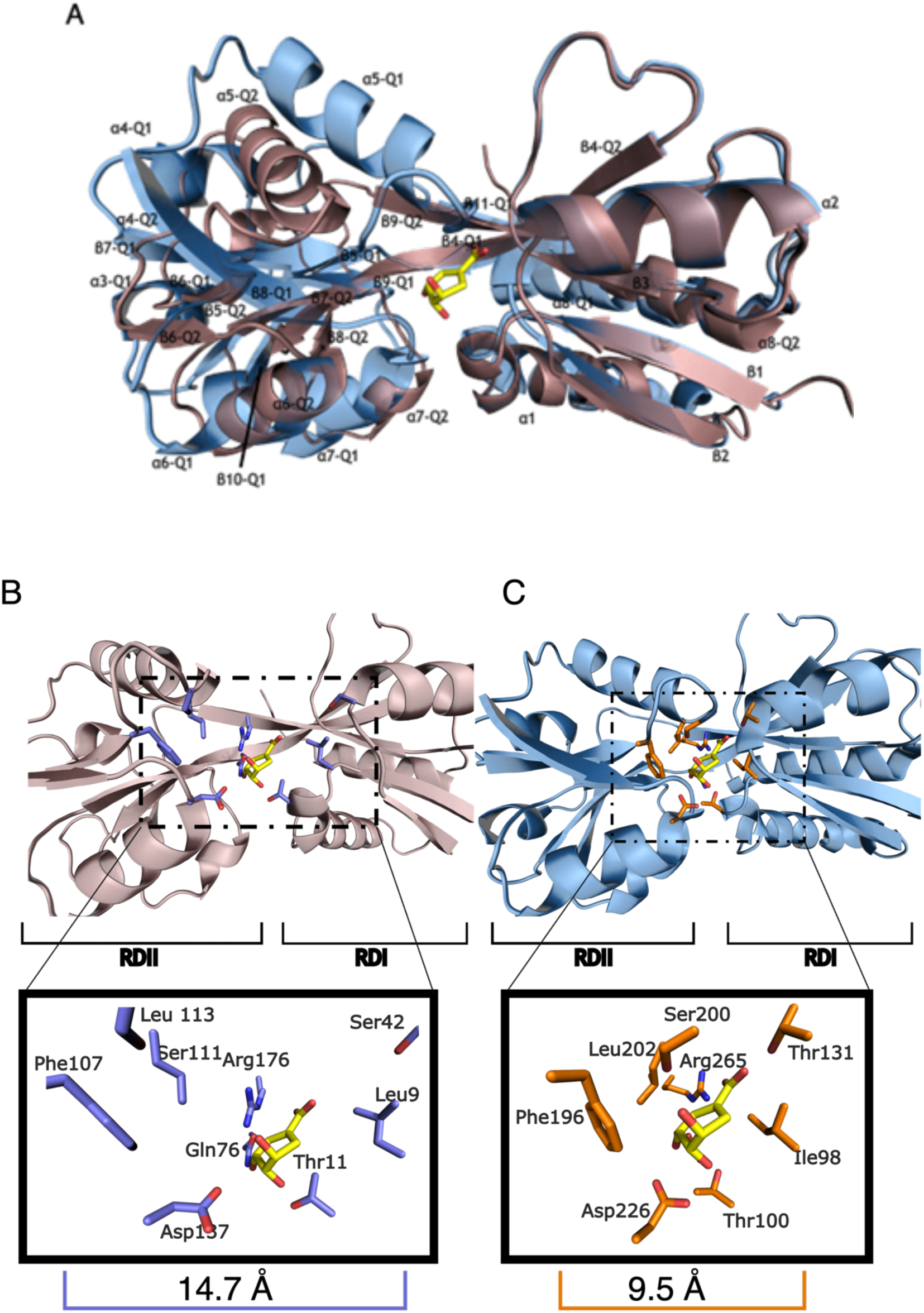
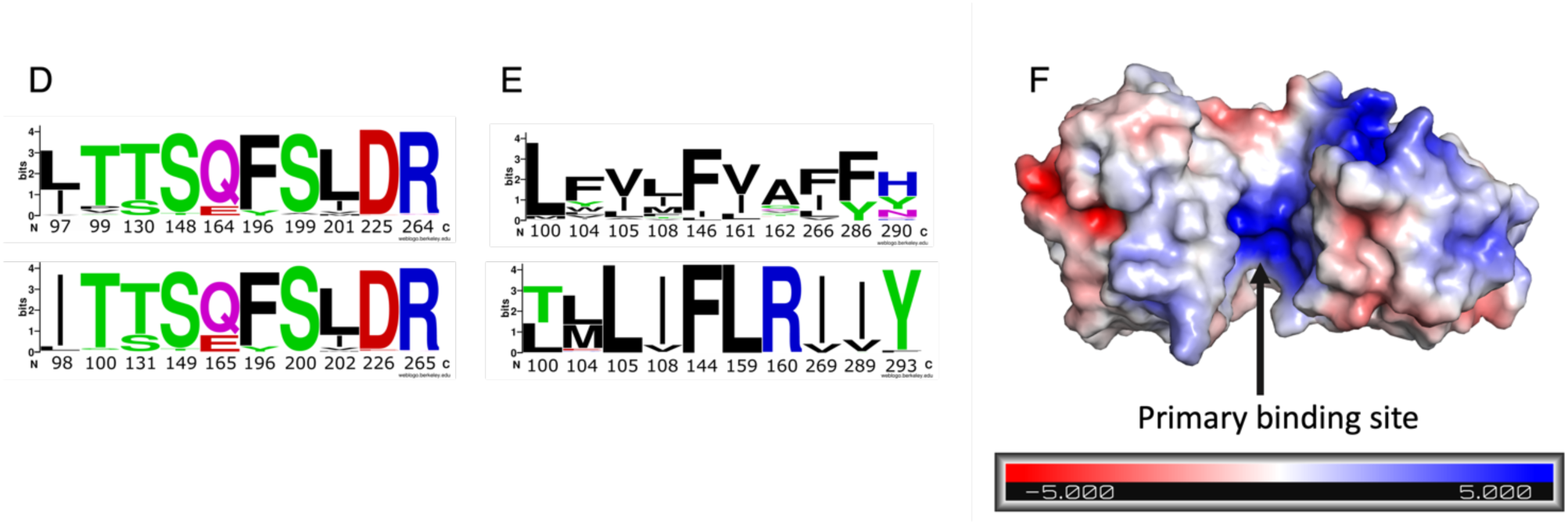
Structural analysis of QuiR2. **A**: Superimposition of QuiR (PDB ID: 5TED; blue) and QuiR2 (brown) cartoons with shikimate from QuiR in yellow. Structural alignment was performed using RDI from both proteins **B**: Cartoon representation of a QuiR2 monomer with proposed ligand binding site residues coloured in blue. Shikimate (yellow) is modeled into the binding site from the QuiR holoprotein. **C:** QuiR (PDB ID: 5TED) cartoon depicting ligand binding site residues in orange. Shikimate (yellow) was co-crystallized in QuiR as part of the study done in Prezioso *et al*. 2018. **D**: Sequence logo of effector binding residues in the QuiR2 (top) and QuiR (bottom) type LTTRs using the Berkeley WebLogo tool. 241 and 173 sequences were aligned for each respective LTTR (Prezioso et al. 2018). **E**: Sequence logo of a putative secondary effector binding site in QuiR2 (top) and of a validated secondary binding site in the BenM LTTR (Ruangprasert *et al*., 2010). 241 and 355 sequences were aligned for QuiR2 and BenM respectively. **F**: Electrostatic surface representation of a QuiR2 monomer. Positive (blue) and negative (red) electrostatic potentials are indicated by a colour gradient. The primary binding site is indicated with an arrow.

**Table 1.** Data collection and refinement statistics. Statistics for the highest-resolution shell are shown in parentheses.

| **Crystal Parameters** | **QuiR2** |
| --- | --- |
| Space group | P 3<sub>1</sub> 2<sub>1</sub> |
| Unit Cell (Å<sup>3</sup>) | 72.659 x 72.659 x 114.17 |
| α, β, γ (°) | 90, 90, 120 |
| **Data Collection statistics** |  |
| Wavelength(Å) | 1 |
| Resolution range | 26.28 - 2.5 (2.589 - 2.5) |
| Total reflections | 25060 (2430) |
| Unique reflections | 12530 (1211) |
| Multiplicity | 2.0 (2.0) |
| Completeness (%) | 99.74 (99.67) |
| Mean I/sigma(I) | 45.60 (4.80) |
| Wilson B-factor | 25.83 |
| R<sub>merge</sub> | 0.0064 (0.131) |
| R<sub>meas</sub> | 0.009051 (0.1853) |
| R<sub>pim</sub> | 0.0064 (0.131) |
| CC1/2 | 1 (0.975) |
| CC* | 1 (0.994) |
| **Refinement and model statistics** |  |
| Reflections used in refinement | 12513 (1211) |
| Reflections used for R<sub>free</sub> | 1269 (122) |
| R<sub>work</sub> | 0.2196 (0.2577) |
| R<sub>free</sub> | 0.2704 (0.2891) |
| No. of non-hydrogen atoms | 1778 |
| No. of macromolecules | 1676 |
| No. of solvent | 102 |
| Protein residues | 208 |
| RMS (bonds) (Å) | 0.006 |
| RMS (angles) (°) | 0.89 |
| Ramachandran favoured (%) | 95.15 |
| Ramachandran allowed (%) | 4.37 |
| Ramachandran outliers (%) | 0.49 |
| Rotamer outliers (%) | 4.89 |
| Clashscore | 8.25 |
| Average B-factor (Å<sup>2</sup>) | 35.37 |
| Macromolecules | 35.53 |
| Solvent | 32.79 |
| **PDB code** | **9O0T** |

QuiR2 structural homologs were determined via a PDB search using the Dali homology server. QuiR was identified as the closest structural homolog (z-score: 22.7, RMSD: 3.7, sequence identity: 44%). The QuiR EBD was structurally aligned with QuiR2 EBD to identify the primary effector binding site. Superimposition of the C-alpha backbone between QuiR and QuiR2 reveals a high degree of structural conservation between the two proteins (**Figure 3A, B**). Based on this analysis, we expect that the primary effector binding region of QuiR2 is located between the two subdomains of the monomeric protein. The amino acid composition for this identified ligand binding site includes the following residues: Leu97, Thr99, Ser130, Ser148, Gln164, Phe195, Ser199, Leu201, Asp225, and Arg264. Sequence logo analysis reveals that these residues are highly conserved in the QuiR2 effector binding pocket (**Figure** 3**C**). The primary binding site of QuiR2 is notably larger than the primary binding site of QuiR; the dimensions of the QuiR2 binding pocket is estimated to be approximately 15 Å by 12 Å, in comparison to the 11 Å by 9 Å for the QuiR ligand binding site (Prezioso *et al.,* 2018).

Some LTTRs have a secondary effector molecule, we therefore analyzed the QuiR2 crystal structure for a secondary binding site (Ezezika et al., 2006). This analysis was conducted with the BenM EBD (PDB ID: 2F78) because they have a well-defined and characterized secondary effector binding site. A position on QuiR2 EBD was identified to superpose with the BenM secondary site (z-score: 18.5, RMSD: 3.4, percent sequence identity: 19). Further, the amino acid composition of this identified site in QuiR2 share high sequence similarity to the BenM secondary site (data not shown). This BenM structure contains the secondary ligand binding site that is also seen on QuiR2. This cavity is flanked by a number of highly conserved hydrophobic residues in QuiR2, as displayed in the globular representation (**Figure** 3**F**). Based on this analysis, the putative QuiR2 secondary binding site is composed of non-polar residues, including Leu100, Leu104, Ile105, Leu108, Ile146, Ile161, Ala162, Phe266, Phe286, in addition to a polar residue, His290. The identified QuiR2 and BenM secondary binding sites share similar residues, most of which consist of Ile and Leu. Interestingly, the position of Ile146 and Phe266 on QuiR2 is switched in the BenM structure. However, Leu104 gates the putative secondary binding pocket of QuiR2, likely regulating the access of the effector molecule to this site. Moreover, this secondary binding site in QuiR2 displayed reduced amino acid sequence conservation as compared to its primary binding site (**Figure** 3**E**). The electrostatic surface potential of QuiR2 shows this secondary ligand binding site is enriched in positively charged amino acids (**Figure 3F**). The overall physiochemical properties of this site together with conservation with the BenM secondary binding site are highly suggestive that an effector molecule can bind to this site on QuiR2.

### Sequence analysis of QuiR2 active site residues

QuiR2 effector binding sites was investigated by sequence analysis to identify conservation of amino acids within characterized binding sites of other LTTRs. Multiple sequence alignment was conducted with QuiR2’s effector binding domain (EBD) and functionally characterized LTTRs including the AmpR, BenM, and QuiR groups of transcriptional regulators (**Figure 4**). This analysis compared amino acid residues that are known to participate in effector binding amongst this group of characterized LTTRs. A total of ten ligand binding residues are highlighted in the QuiR2 sequence. Eight out of ten putative ligand binding residues in QuiR2 are conserved amongst the QuiR group of regulators. In contrast, only six of eight ligand binding residues of BenM and two of eight AmpR residues are shared with the group of ten ligand binding residues conserved between QuiR and QuiR2.

**Figure 4:**
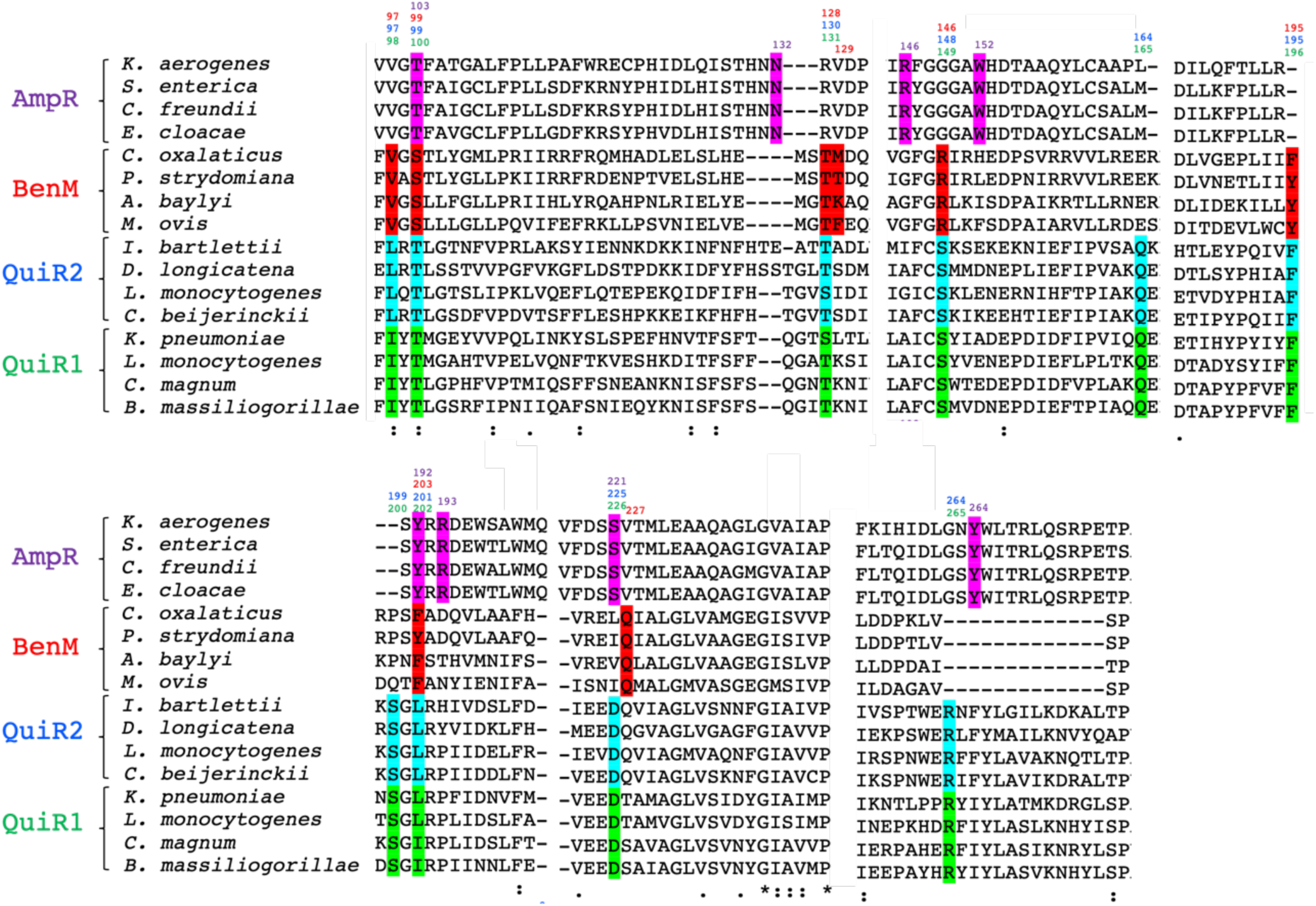
Multiple Sequence Alignment Analysis of QuiR2. Amino acid sequence alignments of QuiR2 against 3 distinct families of structurally characterized LTTR regulators. Represented sequences include the effector binding domain of AmpR, BenM, QuiR2, and QuiR1. Sequences were aligned using the ClustalW Omega software with character counts. Columns with the “.” symbol at the bottom indicates residues that exhibit weak similarity, while the “:” and “*” symbols indicate highly similar and invariant. The active site residues have been highlighted for each corresponding LTTR. Above each colour indicates the position of the residue in the protein. A more detailed sequence alignment is located in the **Figure S3**, along with the organism’s name and NCBI accession number in **Table S1**.

A comparison of the physiochemical properties of the residues that form the ligand binding site indicated that while eight of the ten amino acid residues are conserved between QuiR and QuiR2 in *L. monocytogenes*, the differing two residues show similar chemical properties. This can be seen from Leu97 in QuiR2 which aligned with Ile98 from QuiR and similarly a Ser at position 130 of QuiR2 aligns with a Thr131 from QuiR. The biochemical properties of ligand binding residues in QuiR2 are poorly conserved with BenM and even less with AmpR. A comparison of key residues in the effector binding domain, indicates that QuiR2 shares common features with QuiR in contrast to the other LTTRs.

### Autodock analysis of QuiR2

Using the Dali Server, an initial pairwise structural comparison of QuiR2 with the QuiR EBD was performed. The two proteins superimposed with an RMSD value of 3.6 Å and a Z-score of 27.7 (**Figure 5A**). Although the superimposition of the two structures revealed a high degree of structural conservation, it was evident that the distance between the two structural domains that produces the QuiR2 ligand binding pocket was significantly larger compared to QuiR. Since LTTRs have been experimentally shown to undergo a large conformational shift following effector binding, coupled with the high degree of structural similarity, we reasoned that QuiR2 adopts a similar conformation to QuiR upon ligand binding (Monferrer *et al*., 2010). Hence, to identify a list of candidate ligands, the RDI and RDII domains were independently superposed onto the respective structural domain on QuiR and their relative position was used to generate a QuiR2 conformationally shifted model (**Figure 5A**). Pairwise comparison between the initial QuiR2 structure and the shifted model indicated that the two models are related by an RMSD value of 1.3 Å and a Z-score of 27.8 (**Figure 5B, C**). We expected that this shifted model would represent the ligand bound state of the protein.

**Figure 5:**
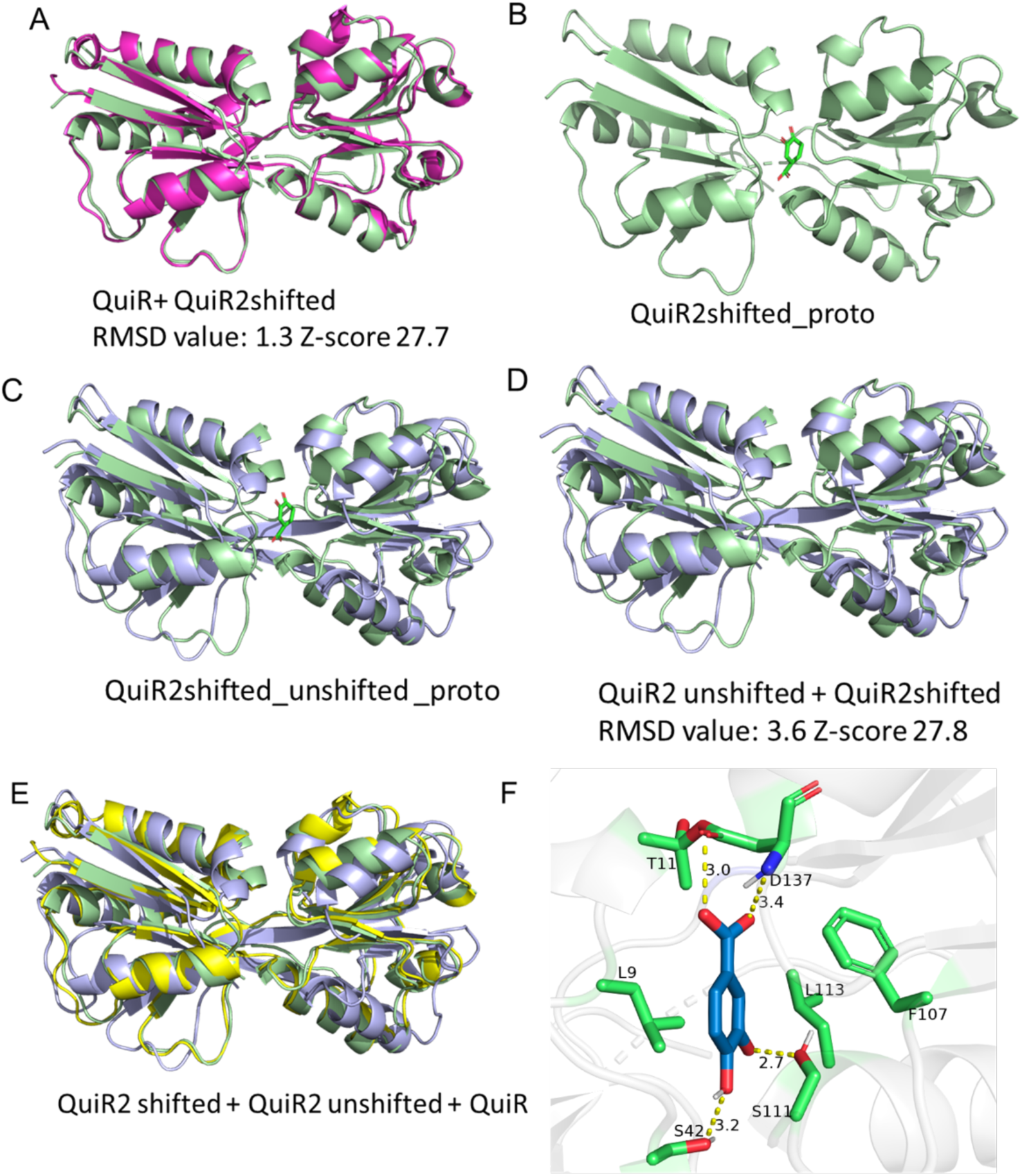
QuiR and QuiR2 Autodock analysis. **A**: Superimposed QuiR and QuiR2 EBD where QuiR2 RDI and RDII were independently superimposed with QuiR. **B**: Shifted QuiR2 EBD as determined from panel A with protocatechuate modeled into its effector binding site. **C**: A comparison of the manually shifted RDI and RDII QuiR2 EBD with unshifted QuiR2 EBD with protocatechuate modeled into the agonist site. **D**: A comparison of the manually shifted RDI and RDII QuiR2 EBD with unshifted QuiR2 EBD without ligands. **E**: Superimposition of QuiR EBD, QuiR2 EBD and QuiR2 EBD with RDI and RDII shifted to better match QuiR. **F**: Modeled protocatechuate binding in the effector binding site of the shifted QuiR2 EBD structure.

From the virtual screen, a list of the top 20 compounds and their corresponding affinity and RMSD values were identified (**Table S2**). This list of candidate ligands includes the following: 9 aromatic compounds, 10 cyclical, and 1 linear. All compounds except for toluol contained hydroxyl groups, followed by carbonyls, or ester functional group. Protocatechuate was identified from the virtual screen with an affinity value of -4.3 kcal/mol. Since protocatechuate is a promising candidate, we docked it into the QuiR2 primary binding site residues participating in this interaction. Residues 4 Å from protocatechuate were selected and displayed in the ligand binding site (**Figure 5F**). Additionally, potential polar contacts and their corresponding distances have been annotated in the ligand binding site. Thr11, Gln10, Leu9, Ser42, and Ser111 were all identified to interact with the modeled protocatechuate.

### QuiR2 effector binding analysis

Differential scanning fluorimetry (DSF) was conducted to identify the QuiR2 effector molecule. Instances where ligand binding influences a protein’s conformation, a shift in the protein melting temperature will be observed with protein-ligand complexation. Since LTTRs are typically regulated by metabolic intermediates or precursor metabolites of the pathway in which they regulate, protocatechuate pathway intermediates were chosen as the primary candidates for our investigation (Maddocks and Oyston, 2008; Monferrer *et al*., 2010). We therefore analysed the change in QuiR2 melting temperatures in the presence of these compounds including protocatechuate. Specifically, we investigated the effect of shikimate, quinate, protocatechuate, dehydroshikimate, formate, chlorogenic acid, caffeic acid, toluene, phenol, magnesium chloride, and tyrosine on QuiR2 melting temperature. A large negative change was observed in the melting temperature with protocatechuate (**Figure 6A**). 10 mM protocatechuate imparted an 8.0℃ reduction (8.0℃; p<0.0001) in QuiR2 melting temperature. Additionally, increasing concentrations of protocatechuate is correlated with a logarithmic change in melting temperature (**Figure 6B**). The resulting denaturation curve obtained with varying concentration of protocatechuate displays a concentration dependent effect on the melting temperature. The crystal structure of QuiR2 revealed two, a primary and a secondary, ligand binding sites. The co-ordinated binding of ligands to both sites could have a distinct role on the biological properties of this regulator.

**Figure 6:**
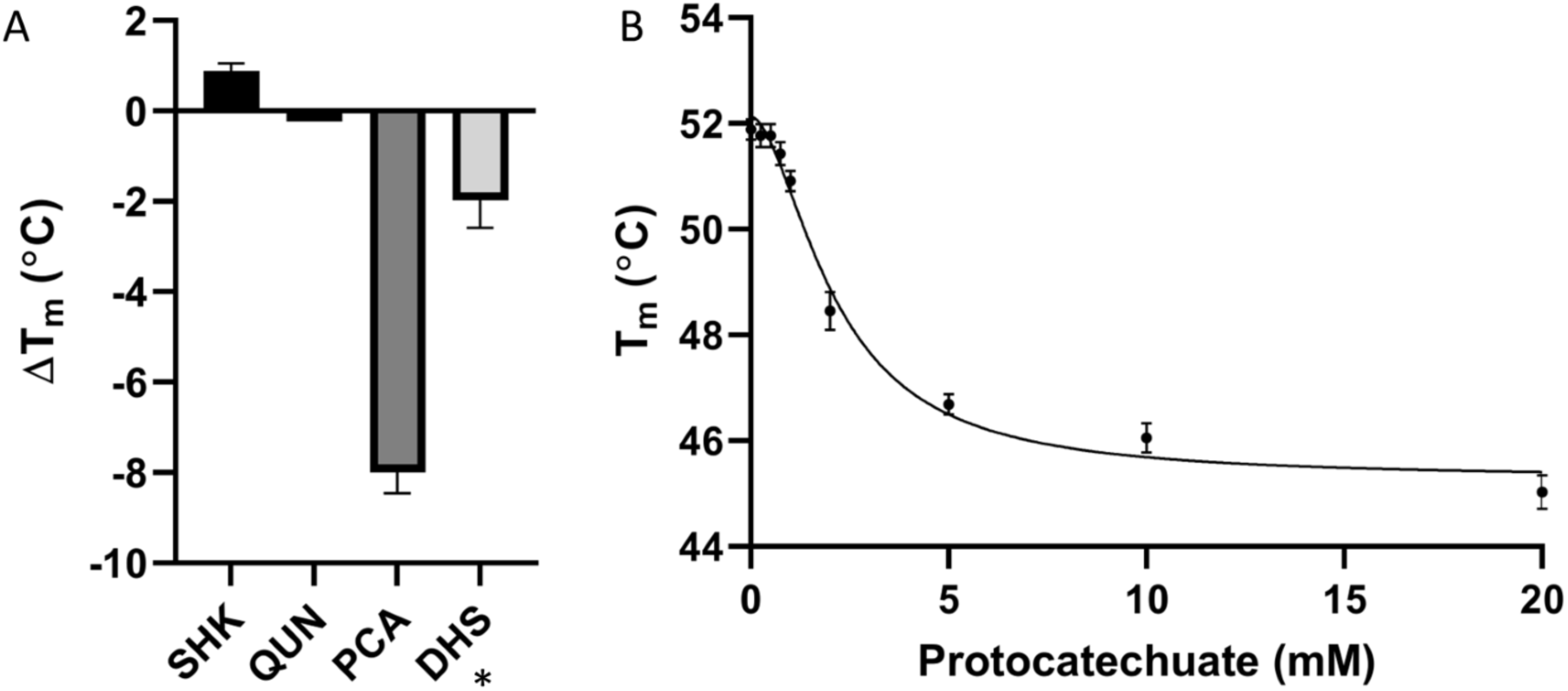
Thermal denaturation analysis of candidate QuiR2 effectors. **A**: A thermal denaturation curve to screen QuiR2 ligands. The differences in melting temperature between the control (no ligand) condition and the shikimate (SHK), quinate (QUN), protocatechuate (PCA) and dehydroshikimate (DHS) conditions are represented as a bar graph. 10 mM of each ligand was added per reaction. Capped error bars represent the standard deviation between 5 replicates (* marked ligand screen was performed in triplicates). **B**: Protocatechuate saturation curve of QuiR2. Capped error bars represent the standard deviation between 7 replicates.

### QuiR2 DNA binding assay

EMSAs were performed to identify the QuiR2 protein DNA binding region(s). As *quiR2* is encoded divergently upstream of *qui2*, the intergenic region of *qui2* was selected for investigation. Furthermore, as the QuiR protein was previously shown to bind to both the *qui1* and *qui2* intergenic regions, we investigated if QuiR2 shares similar binding characteristics. As such we conducted binding assay with QuiR2 and the *qui1* intergenic region (Prezioso et al., 2018). EMSA based analyses reveal that QuiR2 binds both *qui1*’s and *qui2*’s intergenic regions with differing binding affinities (**Figure 7A, B**). Furthermore, when QuiR2 was incubated with 5 mM protocatechuate the mobility of a major fraction of the *qui2* probes increased. This effect was not clearly observed with *qui1* as a higher proportion of the probe was trapped at the top of the gel. Moreover, at lower concentrations of QuiR2 and protocatechuate, *qui1’s* mobility was not clearly influenced by protocatechuate supplementation. Competition assays show that QuiR2 binding to the biotinylated *qui2* probe was completely ablated when the protein was preincubated with 1000 ng (50x the labeled probe) of unlabeled probe thus indicating that this interaction is specific to the *qui2* intragenic region (**Figure 7E**).

**Figure 7:**
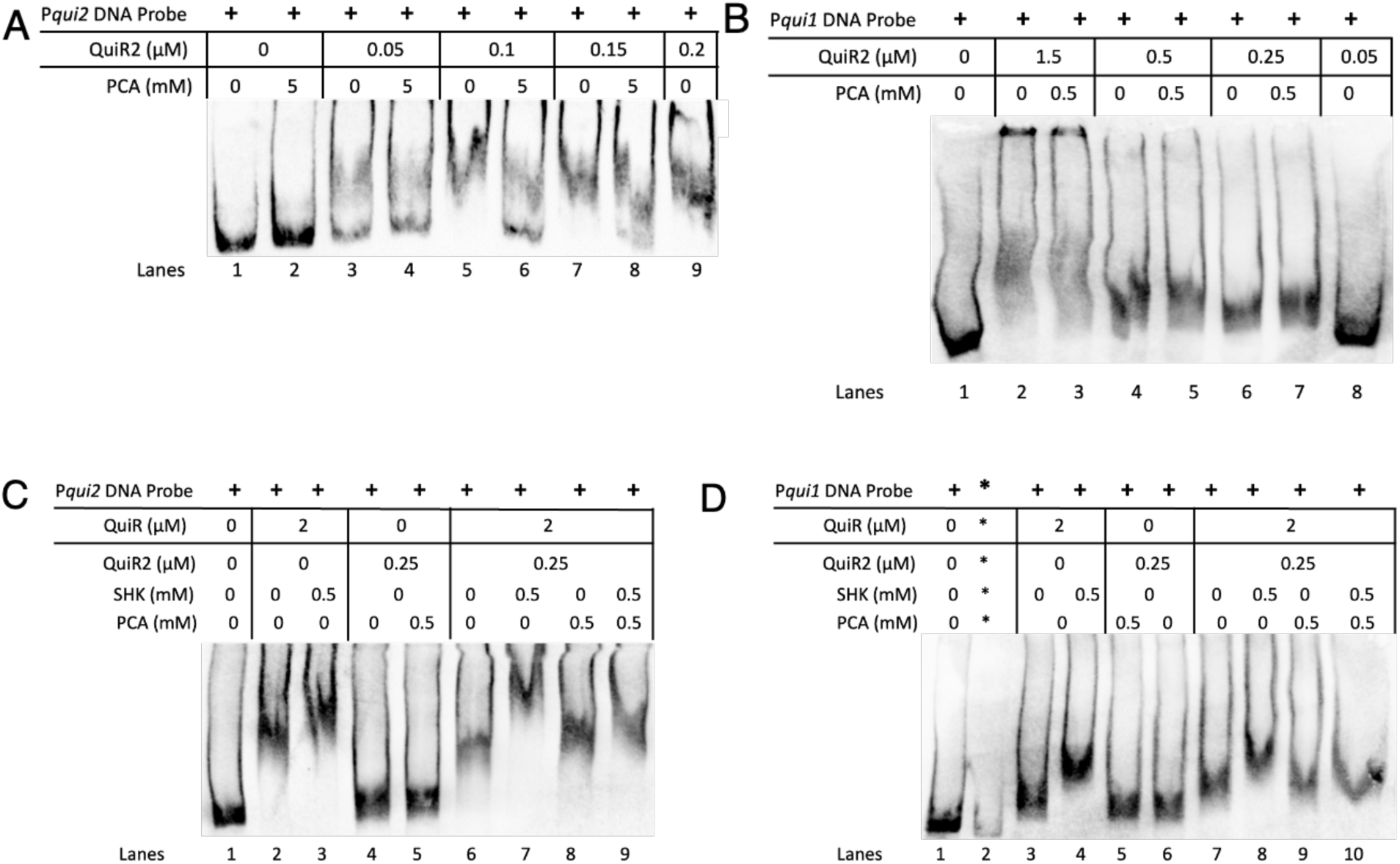
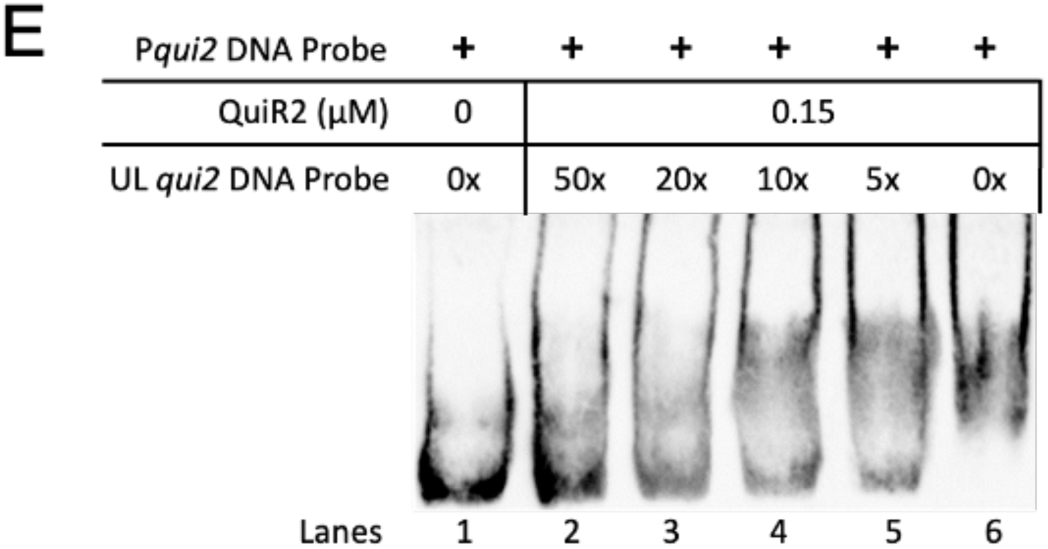
Electrophoretic mobility shift assays for binding of QuiR2 to the *qui1* and *qui2* intergenic regions. All binding reactions contained 20 ng of biotinylated probe. **A**: A gradient EMSA was performed to determine the concentration of QuiR2 protein to bind the *qui2* probe. Reactions were incubated with 0 or 5 mM protocatechuate. **B**: A gradient EMSA was performed to determine the binding of QuiR2 to the qui1 probe. Binding reactions contained 0 or 0.5 mM protocatechuate. **C**: Duo protein incubation of *qui2* probes to assess synergy between regulators. SHK = shikimate. PCA = protocatechuate **D**: Duo protein incubation of *qui1* probes to assess synergy between regulators. **E**: Competition assay to determine the specificity of QuiR2 to qui2’s intergenic region. QuiR2 concentration was kept at 0.15 μM of QuiR2 for samples 2-6. From lanes 2-6 QuiR2 was incubated with decreasing concentrations of unlabeled qui2 probe prior to incubation with biotinylated probe.

Nucleotide sequence analysis indicated that the *qui2* intergenic region is 101 base pairs longer than *qui1*. We posited that the longer *qui2* intergenic region is required to facilitate the concurrent binding of two transcriptional regulators (Figure S3). In order to test this hypothesis, we incubated both *qui2* and *qui1* with semi-saturating concentrations of both transcriptional regulators in the absence of effector ligand and found that this condition induced a mobility shift that is similar to the addition of only QuiR (**Figure 7C,D**). Further, a combination of both transcriptional regulators with shikimate impeded the mobility of the *qui2* DNA probes more so than the QuiR-shikimate treatment. This effect was not observed with the *qui1* intergenic region. Moreover, in both EMSA studies, addition of protocatechuate increases mobility of either DNA probes when both regulators are incubated with shikimate. Notably, the addition of both regulators to *qui2* has a more pronounced impact on the probe motility, reduced mobility, compared to the *qui1* probe. Thus, the longer *qui2* intergenic region and the combined effect of both regulators on the DNA probe mobility would indicate that gene expression from the *qui2* operon is modulated by the action of both regulators together.

### Gene expression analysis by β -galactosidase assay

To further investigate the regulatory function of QuiR2, gene expression analyses were conducted through a ꞵ-galactosidase reporter system. A modified version of the ꞵ-galactosidase assay was performed, where cells were grown without treatments of an inducing agent and at log phase the cells were treated for one hour with co-inducers. The results of this modified protocol produced a slightly higher Miller Unit value in all samples, though it allows for analysis with potentially toxic compounds because the cells grow to sufficiently high density prior to supplementation with these compounds (**Figure S**4). Reporter constructs were designed to evaluate the role of QuiR2 on both *qui1* and *qui2* operon gene expression and also QuiR2 autoregulation as shown in Figure 8. The expression of LacZ under the *qui1* promoter and QuiR regulator in *B. subtilis* was used as a positive control (Prezioso *et al*., 2018). Next, we overexpressed QuiR2 to test its regulatory effect on both *qui1* and *qui2* operons and observed that the protein has a strong repressive effect on both operons. The strain containing xylose-inducible *quiR2* gene and *lacZ* gene inserted at the beginning of *qui2* operon did not produce any detectable LacZ activity even with the addition of xylose. Further, the addition of protocatechuate did not alleviate the repressive activity of QuiR2. Similarly, the placement of LacZ at the beginning of *qui2* operon and natively expressed QuiR1 did not alleviate the repressive activity of QuiR2. Even with basal level of QuiR2, without the addition of xylose, the level of LacZ activity was close to background activity. Interestingly, the addition of shikimate in this non-induced condition of QuiR2 resulted in a measurable level of LacZ activity. Thus, indicating that shikimate activated QuiR overcame the repressive action of QuiR2. This is further demonstrated in the condition in which xylose was used to induce the expression of QuiR2. Under this condition, the presence of shikimate activated QuiR was insufficient to counteract QuiR2 repression.

**Figure 8:**
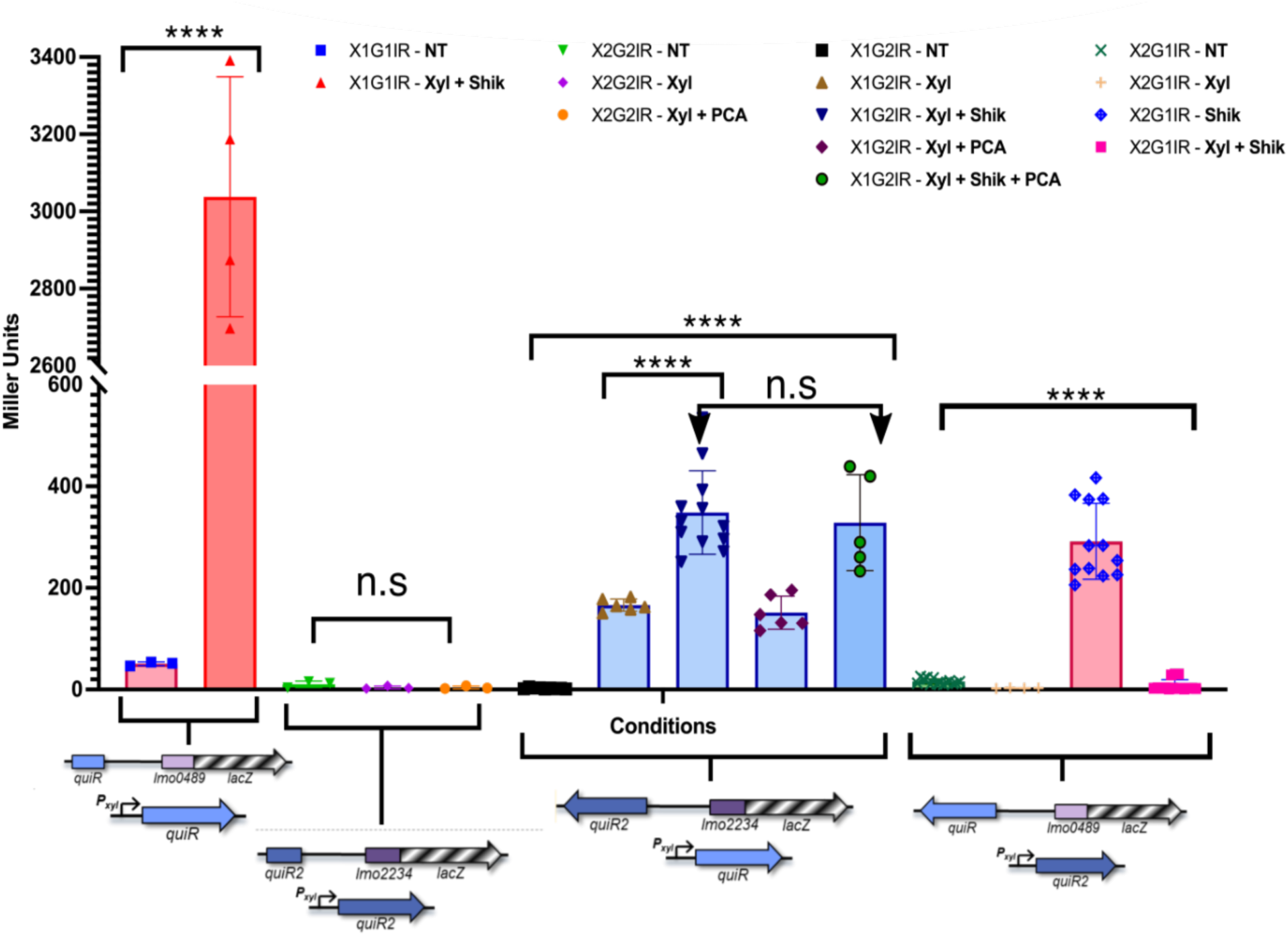
β -galactosidase Assay. Gene expression analyses of *qui1* and *qui2* operons in a *B. subtilis* background. Genetic constructs are visually described with graphics underneath each data bar. For naming convention, *B. subtilis* was transformed with pAX01: QuiR (X1), pAX01:QuiR2 (X2), pDG1661: *quiR*+*qui1* intergenic region (G1IR), pDG1661:*quiR2+qui2* intergenic region (G2IR). NT= no treatment, Xyl = 0.5% Xylose, Shik = 1 mM shikimate, PCA = 1 mM protocatechuate. All assays were conducted without delayed induction. Replicate counts reflected by data points.

To investigate if there is concomitant regulation by QuiR and QuiR2, we assayed *B. subtilis* strains with both regulators. In this scenario, QuiR is natively expressed and QuiR2 expression is under a xylose-inducible promoter and the *lacZ* gene is inserted at the beginning of the *qui2* operon. In the absence of xylose, the basal level of QuiR was insufficient to overcome QuiR2 repressive action. However, induction of QuiR expression with xylose overcame this repression as observed with the increased LacZ activity. Addition of shikimate to the xylose induced condition resulted in further increased in LacZ activity. However, the addition of protocatechuate to the xylose induced condition did not have any effect on the level of LacZ compared to the untreated xylose induced condition. Surprisingly, the addition of both shikimate and protocatechuate effectors, showed an increased in LacZ activity thus indicating that the shikimate activated QuiR is sufficient to offset the QuiR2 repressive activity. These findings overall indicated that QuiR2 is functioning as a transcriptional repressor and this can be overcome by shikimate-bound QuiR. It was shown that shikimate binds much stronger to QuiR compared to protocatechuate and as such is consistent with our observation when both effectors were used in the study above (Prezioso *et al*., 2018).

## Discussion

### QuiR-type LTTRs regulate atypical protocatechuate metabolism

The biochemical function of the dehydroshikimate dehydratase (DSD), QuiC2, has been implicated in the biosynthesis of protocatechuate in *L. monocytogenes’* (Xue *et al*. 2020). However, the biological role of this compound in *L. monocytogenes* remains unclear. To gain an understanding of the biological role of protocatechuate to *L. monocytogenes*, we investigated the regulatory mechanism of two transcriptional regulators, QuiR and QuiR2, that are involved in modulating gene expression for proteins in the protocatechuate biosynthetic pathway (Prezioso *et al*. 2018). In order to understand the functional relationship between the two QuiR regulators and other LTTRs, they were aligned against other characterized LTTRs and phylogenetics analysis was conducted. This approach allowed us to establish that QuiR and QuiR2 form distinct phylogenetic clusters from each other and from other LTTRs. Since the functionally characterized LTTRs formed independent clusters, we envisioned that both QuiR and QuiR2 clusters represent distinct molecular functions. Moreover, QuiR and QuiR2 diverge from other LTTRs involved in the metabolism of aromatic compounds such as BenM, CatM, and AaeR (Van Dyk *et al*., 2004; Ezezika *et al*., 2006). The latter regulators are associated with the *β*-ketoadipate pathway, and their distinct clustering reflects the absence of this pathway in *Listeria*. Similarly, previous phylogenetic analyses revealed that QuiC2 also diverges from established functional groups of DSDs, further suggesting that enzymes from this pathway are involved in novel biological processes (Xue *et al*. 2020).

### Regulatory role of QuiR2

QuiR is a shikimate-dependent transcription factor which regulates *qui1* and *qui2* operons gene expression, both of which encodes proteins for protocatechuate biosynthesis (Prezioso et al., 2018). Characterization of the regulatory role of QuiR was conducted by measuring *qui1* and *qui2* operon gene expression through β-galactosidase assays in the absence of QuiR2 interaction (**Figure** 8). EMSA was used to established that QuiR binds to *qui1* and *qui2’*s intergenic regions to regulate gene expression from both operons (**Figure** 7). In this work, studies with the transcriptional regulator QuiR2, for *qui1* and *qui2* operons, was undertaken together with QuiR to further elucidate the regulation of protocatechuate biosynthesis. Through EMSA experiments, it was established that, similar to QuiR, QuiR2 binds to *qui2* intergenic region to regulate gene expression. Further, it was shown that both QuiR and QuiR2 can bind concurrently to *qui2* intergenic region. The binding of both regulators to this intergenic region is consistent with the LacZ reporter assay which indicate that QuiR2 is functioning as a transcriptional repressor. The accommodation of two transcriptional regulators suggests that they function together to regulate protocatechuate metabolism. Indeed, β-galactosidase reporter assays conducted using both regulators, reveal that QuiR2 attenuates the activating effects of QuiR. Compared to the studies by Prezioso *et al*. 2018, the overall expression of the two operons is an order of magnitude lower when QuiR2 was expressed together with QuiR, in contrast to QuiR alone. This suggests that expression levels of *qui1* and *qui2* genes, when QuiR is expressed alone, may not reflect biologically relevant levels of expression as QuiR2 negatively regulates operon expression. Preliminary EMSA analyses suggest that QuiR and QuiR2 bind concomitantly to *qui2* intergenic region. This is consistent with the observation that the presence of both regulators imparts a stronger reduction in mobility of the biotinylated *qui2* DNA probe compared to mobility with individual regulators. In contrast, neither regulator binds to the *qui1* intergenic region in the absence of the effector molecule. This would indicate that the regulation of gene expression from the *qui1* operon is tightly controlled by the accumulation of the respective effector molecules. As shown by EMSA for *qui1* intergenic region, shikimate induces QuiR binding to the *qui1* probe. The addition of protocatechuate to this QuiR-shikimate-*qui1* complex results in disassociation of the regulator from the DNA probe. The repressive action of QuiR2 is also observed with the QuiR-*qui1* probe. There seems to be a noticeable increase in mobility of the QuiR-*qui1* probe with the addition of QuiR2 and protocatechuate. Therefore, one can postulate that either the effector or the effector-QuiR2 complex is participating in disrupting QuiR from the *qui1* probe.

Interestingly, the findings from our DSF study suggests that protocatechuate is a ligand to QuiR2 as evidenced by the protocatechuate concentration dependent change in QuiR2’s melting temperature. In this analysis, protocatechuate induces a conformational reorganization of QuiR2 which is correlated with a reduction in the melting temperature of the protein. Similarly, this effect was observed with DNA binding studies using the *qui2* intragenic region. Interestingly, we also observed similar effect of protocatechuate on QuiR-*qui1* probe interaction. Further, protocatechuate supplementation in our LacZ reporter assays with QuiR and QuiR2 had a negative effect on gene expression for both *qui1* and *qui2* operons. Interestingly, the LacZ assay indicated the addition of shikimate overcame this destabilizing effect of protocatechuate on QuiR activity. Based on these findings, it is expected that QuiR2 is functioning as a transcriptional repressor and the binding of protocatechuate relieves this repressive effect by destabilizing the protein interaction to the *qui2* promoter. Similar regulatory mechanism of gene expression has been frequently observed in microbial systems (Terán et al., 2006; Sakono and Hayakawa, 2021).

### Biological role of QuiR2 in *Listeria innocua*

In this study, we have investigated the role of two transcriptional regulators, QuiR and QuiR2 on modulating gene expression from two genomic operons, *qui1* and *qui2*. It has been established that genes from both operons encode proteins that collectively are involved in redirecting shikimate to produce protocatechuate (Prezioso et al., 2018; Xue *et al*. 2020). The role of protocatechuic acid in *Listeria* species have been elusive because these microorganisms do not have the metabolic capacity to utilize this compound. However, it was proposed that protocatechuate can be released into the environment and allows *Listeria* species to establish synergistic relationship with other organisms. Since shikimate is a vital metabolic intermediate for key aromatic compounds, including aromatic amino acids, in microorganisms it is of utmost importance to finely tune its distribution for the synthesis of these compounds. Instances when shikimate accumulates in excess, it is redirected to protocatechuate to facilitate microbial interaction. In this study, we established that both transcriptional regulators, QuiR and QuiR2, play crucial regulatory roles in establishing the distribution of shikimate to aromatic compounds and protocatechuate synthesis. We developed a model in which QuiR functions as a transcriptional activator, QuiR2 a repressor, and shikimate as a key modulator for QuiR activity. As shown, QuiR function is negatively regulated at high protocatechuate concentration, and this effect is circumvented by addition of shikimate. Therefore, in a biological context it is expected that QuiR2 binds to the *qui2* promoter, preventing elevated gene expression from this operon and thus allowing shikimate to be used for vital aromatic compounds biosynthesis. As shikimate accumulation increases, it activates QuiR which binds to both *qui1* and *qui2* operons and resulted in increased gene expression from both operons and this redirects shikimate to protocatechuate synthesis. Once high level of protocatechuate accumulates in the organism, it activates the repressive activity of QuiR2 to prevent QuiR from continued gene expression from the respective operons. This reduces the level of shikimate being directed to protocatechuate production. Overall, this study builds on previous observations, which are changing the long-standing paradigm that the shikimate pathway exclusively leads to the biosynthesis of aromatic compounds, including the three aromatic amino acids. The shikimate pathway plays an important role in directing shikimate and quinate to produce protocatechuate which can be used to facilitate microbial interactions in addition it can be used as an important energy producing compounds via the *β*-ketoadipate pathway.

**Figure 9:**
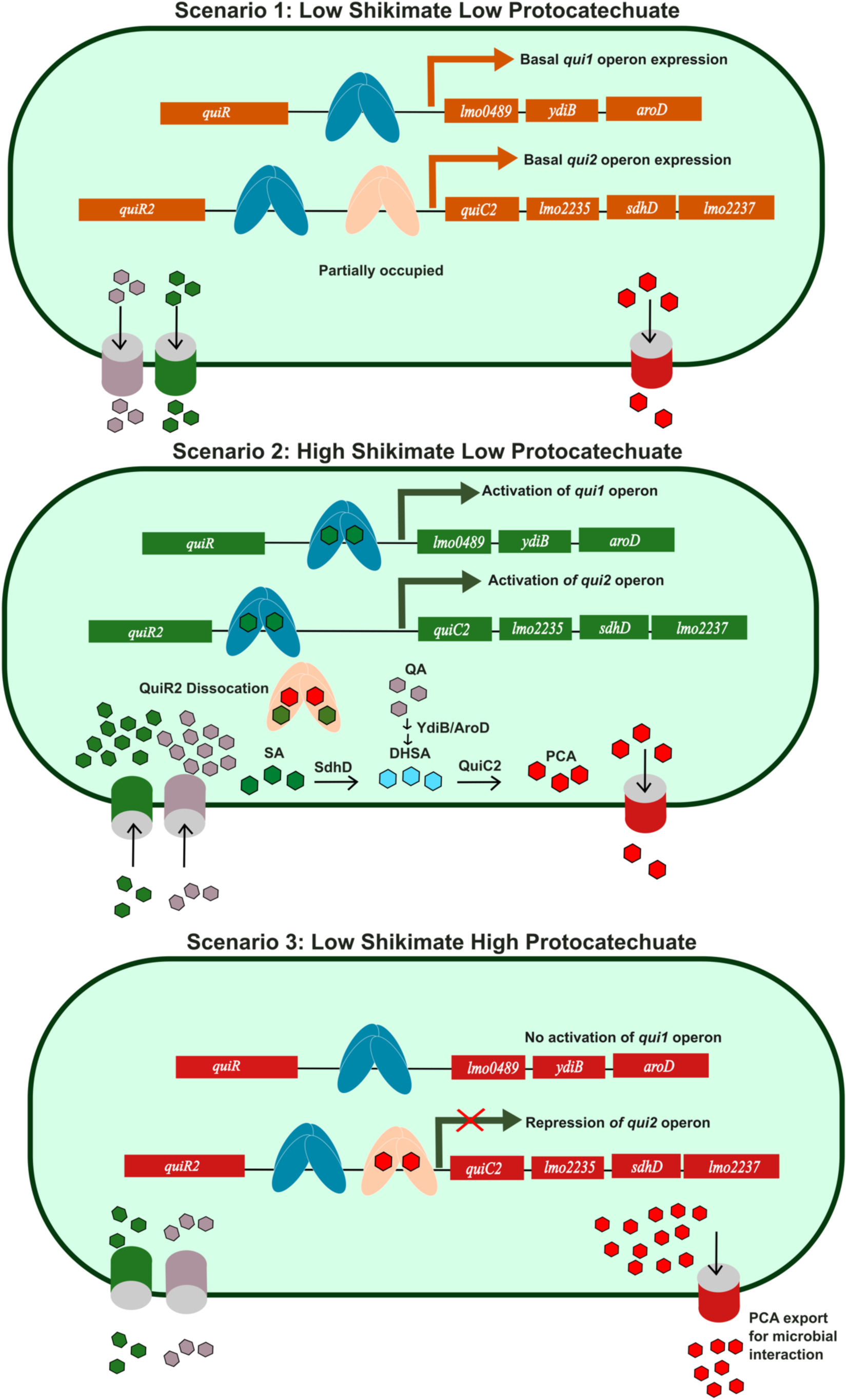
Newly proposed regulatory model for protocatechuate metabolism in *Listeria* species. Regulatory model for protocatechuate metabolism in *Listeria* species considering cases of low and high shikimate availability. Green boxes represent *Listeria* cells. Mauve hexagons= quinate, green hexagons= shikimate, red hexagons= protocatechuate, mauve cylinder= quinate transporter, green cylinder= shikimate transporter, red cylinder= protocatechuate cylinder. Teal ovals= positive regulator, QuiR, peach ovals= negative regulator, QuiR2. Arrows indicate multistep reactions and metabolite movement through transporters. In conditions of low shikimate, *Listeria* conserves shikimate for chorismate and AAA biosynthesis because of QuiR2 downregulating expression of *qui1* and *qui2* genes. In high shikimate conditions, carbon is committed to protocatechuate production as QuiR induces expression of *qui1* and *qui2* genes. Due to size differences in the intergenic regions of *qui1* and *qui2*, QuiR may competitively displace QuiR2 from the *qui1* promoter, whereas QuiR and QuiR2 may concomitantly bind *qui2*’s intergenic region. Under high protocatechuate accumulation, QuiR2 represses *qui2* operon and protocatechuate is exported to facilitate microbial interaction.

## Material and Methods

### Bacterial strains, plasmids, and growth conditions

Bacterial strains and plasmids are listed in **Table S3**. *Escherichia coli* strains were grown in Luria Bertani (LB) media. *E. coli* strains carrying the pET28a and pET28mod plasmids were selected using 50 μg ml^−1^ of kanamycin. *E. coli* strains carrying the pLSV101 plasmid were selected using 300 μg ml^−1^ of erythromycin at 30 ℃ (Joseph et al., 2006). *L. innocua* Clip11262 was grown in bovine heart infusion media (BD BactoTM). All cells were grown at 37 ℃ while shaking (200 rpm) unless stated otherwise. *L. innocua* transformants carrying pLSV101 were grown with 5 μg ml^−1^ of erythromycin. For *quiR2* (*lmo2233*, GenBank ID: CAD00311.1) cloning, *Listeria monocytogenes* EGD-e genomic DNA was kindly provided by Dr. Nelly Leung from the Department of Molecular Genetics at the University of Toronto.

### Protein expression and purification

*E. coli* BL21 cells carrying plasmid pET28MOD:*quiR2,* pET28MOD:*quiR2*Δ*87* or pET28MOD:*quiR2*Δ*89*, respectively, were inoculated in LB broth with 50 µg ml^−1^ kanamycin at 37 ℃ until the cultures reached logarithmic phase (OD_600_ 0.4 to 0.8). Protein expression was induced with 0.4 mM isopropyl-*β*-thiogalactopyranoside (IPTG) and cultures were incubated at 18℃ for 16 hours. Cells were centrifuged with an average of 4790 x*g* for 15 minutes and the resulting pellet was resuspended in 30 mL of binding buffer (50 mM TRIS-HCl, 5 mM imidazole, 5% glycerol, 500 mM NaCl). Protease inhibitor was added and the cells were lysed via French pressure cell press, followed by sonication with the following settings: 80 duty, 5 output, 30 seconds on and 30 seconds off for 5 cycles.

Following centrifugation, the soluble fraction was purified via Ni-NTA affinity chromatography as described in Peek *et al*. 2016. Briefly, binding buffer (50 mM TRIS-HCl pH 7.5, 5 mM imidazole, 5% glycerol, 500 mM NaCl), wash buffer (30 mM imidazole) and elution buffer (300 mM imidazole) were used to purify the protein of interest. 1 mM of EDTA and 0.33 mM of dithiothreitol (DTT) were added to the protein samples. Tobacco etch virus (TEV) protease was used to cleave the hexa-histidine tags during overnight dialysis (10 mM TRIS-HCl pH 7.5, 250 mM NaCl, 5% Glycerol, *β*-mercaptoethanol) at 4℃. A second Ni-NTA affinity chromatography was performed in order to purify un-tagged protein. In the second purification, untagged protein is collected in the binding and wash buffers, whereas uncleaved protein is eluted with elution buffer. Subsequently, the protein was dialyzed overnight at 4℃ in the same buffer as shown above without TEV and concentrated by centrifugation with a 10 kDa Millipore concentrator at 2000 x*g*.

Full-length QuiR2 was purified via fast protein liquid chromatography (FPLC). Concentrated QuiR2 fractions were injected into the 16/60 Superdex 200 size exclusion column attached to a AKTA purifier system (GE Healthcare, Uppsala, Sweden). The FPLC column was equilibrated with 50 mM TRIS-HCl pH 7.5, 500 mM NaCl.

### Protease analysis

An *in silico* protease analysis was performed in order to identify proteolytic cleavage of the full length QuiR2. The full length QuiR2 sequence was submitted to ExPASy PeptideCutter to identify potential protease cleavage sites ( The Proteomics Protocols Handbook). The output list was examined to identify proteolytic agents found in the protein purification conditions. Amino acid residue 87 was identified as a position on the protein sequence that is prone to cleavage by hydroxylamine. Thus, we prepared *quiR2*Δ*87* and *quiR2*Δ*89* constructs for effector binding domain studies.

### Virtual Screening Analysis of QuiR2

The structure-based virtual screening analysis was performed using Autodock and its suite of computational tools against a library of ∼13,000 metabolite compounds from the ZINC15 database ( Trott and Olson, 2009). To prepare QuiR2 for the virtual ligand screen, the PDB structure was initially aligned with the QuiR:EBD docked with shikimate (PDB: 5TED). Since LTTRs have been previously described to undergo a conformational shift following ligand binding, we superimposed the crystal structure of QuiR2 with the existing QuiR-shikimate bound to gain insights on the conformational changes that occur upon ligand binding. The resulting QuiR2 structure was used in a virtual ligand screen with Autodock Vina. The Dali protein structure comparison server (http://ekhidna2.biocenter.helsinki.fi/dali/) was used to obtain root mean square deviation (RMSD) values and Z-scores for pairwise comparison of QuiR, QuiR2 shifted, and unshifted structures (Holm, 2019). Subsequently, the graphical user interface Autodock Tools, was used to convert the QuiR2 receptor into a. pdbqt file by removing water molecules, adding polar hydrogens, and computing Gasteiger charges. To identify the location for docking, the QuiR2 structure was superimposed with QuiR and a center grid box with dimensions x= -18.972, y= -0.2578, and z= -2.544 was generated to encompass the shikimate binding site. The area of the grid box was 14 x 16 x 16, equal to 3,584 Å^3^ The virtual ligand screen was conducted using an AutoDock python script generated by Sari Sabban, 2018 and Vina executable. Using the script, the grid box dimensions were specified, and each ligand was subjected to an exhaustiveness of 8 during the screening process. To increase computing efficiency, the Niagara supercomputer at the SciNet HPC Consortium was used ( Loken *et al*., 2010; Ponce *et al*., 2019). Using Vina, each ligand exhaustiveness computed an affinity (kcal/mol) and RMSD value to identify energetically favourable ligands. Once the initial screen was complete, the top 20 ligands were rescreened using an exhaustiveness of 15 to ensure accuracy and reproducibility of the results. The modified QuiR2 structure was docked with the ligand of interest with the lowest affinity value.

### Protein crystallization

The QuiR2 ligand binding domain (residues 89-204) was crystallized using the hanging drop vapour diffusion method. The hanging drops contained 2 µL of purified QuiR2 protein (27.2 mg ml^−1^) and 2 µL crystallization solution supplemented with one of the following ligands: 50x concentrated *L. innocua* metabolite extract, 500x concentrated *L. innocua:ΔDSD* metabolite extract, 2mM dehydroshikimate. The crystallization solution consisted of 0.2 M potassium thiocyanate and 22% polyethylene glycol 3,350. Individual protein crystals obtained were flash frozen in liquid nitrogen. Briefly, protein crystals were initially transferred to the crystallization solution with 7.5% ethylene glycol and 7.5% glycerol which served as the cryosolution. Crystals were rapidly transferred to a Liquid Nitrogen bath and stored for X-ray data collection. X-ray diffraction data were collected using the synchrotron beamline at the Brookhaven National Laboratory (BNL).

### β -galactosidase assays

β -galactosidase assays were prepared and conducted as described in (Prezioso et al., 2018). Briefly, *B. subtilis* 168 were transformed by inducing starvation-related competence. Recombinant pDG1661 transformants were selected using chloramphenicol and screened for their ability to hydrolyze starch. Recombinant pAX01 transformants were selected using erythromycin and screened through PCR. For control strains, empty pAX01 and pDG1661 were employed. *B. subtilis* transformants were grown overnight in LB broth at 37 °C and 200 RPM, and then diluted 1/100 into 5 mL fresh LB media with 0.5% xylose and 1 mM shikimate or protocatechuate. At an OD600 between 0.6-0.8, cells were washed and pelleted and resuspended in working buffer [60 mM Na2HPO4, 40 mM NaH2PO4, 10 mM KCl, 1 mM MgSO4, 20 mM β-mercaptoethanol (pH 7.0)] with 1mg/mL lysozyme and incubated at 37 °C for 45 min. Ortho-nitrophenyl-β-galactoside (ONPG) was added to a final concentration of 0.67 mg/mL, and individual reactions were stopped when a yellow color was apparent by adding 0.3 M Na_2_CO_3_. Reaction start and stop times were recorded and reactions were centrifuged to pellet debris, and the OD_420_ and OD_550_ for each reaction was determined with a Cary 50 BioUV–Vis spectrophotometer.

### Differential scanning fluorimetry

25 µL reactions containing 2 µM of QuiR2Δ89, 5X SYPRO orange protein gel stain (Sigma), and HEPES screening buffer (pH 7.5) were prepared as described by Allali-Hassani *et al*. 2005. 10 mM of the following compounds were added for ligand screening: shikimate, quinate, protocatechuate, dehydroshikimate, formate, chlorogenic acid, caffeic acid, toluene, phenol, magnesium chloride, and tyrosine. DSF ligand screen was also performed with *Arabidopsis thaliana* Col-0 plant root exudate, *L. innocua* metabolite extract, *L. innocua: ΔDSD* metabolite extract, and *L. innocua: ΔquiR2* metabolite extract. Additionally, a protocatechuate saturation curve was generated using a concentration gradient ranging from 0 to 20 µM. Six replicates of each reaction were prepared in a 96-well plate and the thermal shift was performed with a temperature gradient from 25 ℃ to 95℃ at a rate of 1℃ per minute. All reactions were run in a Real-Time PCR Thermocycler (CFX Real-Time System C1000 Thermal Cycler, Bio-Rad) and fluorescent output was recorded at 0.4℃ increments. Thermal shift curves were processed and protein melting temperatures were calculated using the ‘CFX96 Manager’ software. Melting temperatures were plotted and analyzed using the GraphPad Prism 8© software.

### EMSA

EMSA was performed with the purified full-length QuiR2 protein as well as the same biotinylated probes as published in *Prezioso* et al. 2018. Briefly, a 225-nucleotide and a 402-nucleotide biotinylated DNA probe were designed to amplify the intergenic region preceding the *qui1* and *qui2* operons respectively. The *qui1* probe was amplified from *Listeria monocytogenes* EGD-e gDNA using the 5’ end biotin-labeled reverse primer (OR:R-Biotin) and the unlabeled forward primer (OR:F). To produce an unlabeled probe, the OR:R-Biotin reverse primer was substituted with the unlabeled variant (OR:R). The *qui2* probe was similarly amplified from *L. monocytogenes* EDG-e gDNA using the 5’ end biotin-labeled reverse primer (OR2:R-Biotin) and the unlabeled forward primer (OR2:F). The unlabeled *qui2* probe was also produced by substituting the OR2:R-Biotin reverse primer with the unlabeled OR2:R reverse primer. The 20 µL binding mix used for EMSA consisted of a binding buffer without EDTA, 20 ng labeled DNA probe, 25 ng/µL poly (dl-dC) non-specific competitor DNA, and varying concentrations of purified QuiR2 protein and protocatechuate. The binding mix was incubated at room temperature for 30 minutes prior to gel electrophoresis. Following incubation, samples were separated on a 5% non-denaturing polyacrylamide gel made with ice-cold 0.5x Tris-borate EDTA (TBE; pH 7.5) buffer. The gel was run at 70 V for 10 minutes followed by an increase in voltage to 115 V for 60 minutes. DNA transfer to a positively charged nylon membrane was done by semi-dry electro-blotting at 380 mA for 30 minutes in 0.5 TBE. The membrane was treated with 300 nM UV-lightbox for 30 mins to crosslink DNA and then incubated in a blocking buffer (4% milk in TBS-T) overnight at 4°C. Membranes were washed 3 times in TBS-T prior to a 30-minute room temperature-incubation with streptavidin-HRP (Cell Signaling Technology, diluted to 1/5000 in 5% BSA-T). The membranes were then washed in TBS-T again and the DNA was detected with Clarity ECL Western Blotting Substrate (Bio-Rad) and exposure to X-ray film. For competition assays, prior to adding labeled DNA, binding reactions with up to 50x excess unlabeled probes were incubated first for 20 minutes ( Prezioso *et al*., 2018).

### Bioinformatic analyses

#### Multiple sequence alignment

The effector binding domain sequences belonging to AmpR, BenM, QuiR, and QuiR2 families were aligned through the Clustal Omega program **(Table S1).** The alignment was annotated to show the ligand binding residues and their corresponding positions for each distinct class of LTTRs. The EBD amino acid sequences from AmpR (PDB:3KOS), BenM (PDB:2F6G), and QuiR (PDB:5TED) were obtained from the Protein Data Bank (PDB). The corresponding sequences were used as the query sequence in the Basic Local Alignment Search Tool (BLAST) to obtain the top 1000 hits (Altschul *et al*., 1990; Gish and States, 1993). Subsequently, 3 unique species and the initial query sequence were used for alignment for each LTTR.

#### Phylogenetic analyses

LTTRs with various biological functions were used to query for lists of LTTRs. These LTTRs include: BenMs, CatMs, AmpRs, AaeR, QuiRs, and QuiR2s. The top 1000 hits from BLAST for each LTTR group were filtered for non-redundant sequences and included together for a multiple sequence alignment using Clustal Omega. 761 sequences were included in a maximum likelihood phylogenetic analysis using the FastTreeMP program from XSEDE ( Miller A *et al*., 2010). The phylogenetic analysis included 1000 bootstraps using the Jones-Taylor-Thornton (JTT) model and a penalty score of 1. The output from FastTreeMP was processed using the ITOL software and labeled using Inkscape (Letunic and Bork, 2016).

## Supporting information

Supplemental Table 4-Phylogenetic Sequences

## Acknowledgements

This work was supported by a grant to D.C. from the Natural Sciences and Engineering Research Council (NSERC) of Canada (Grant No. RGPIN-2015-06747). We thank Dixon Ng for collecting and processing the crystallography data. This research used resources of the National Synchrotron Light Source II, a U.S. Department of Energy (DOE) Office of Science User Facility operated for the DOE Office of Science by Brookhaven National Laboratory under Contract No. DE-SC0012704.

