## Supplemental Figures 1-4, Supplemental Table 1-3 for "The coordinated regulatory roles of two LysR-Type Transcriptional Regulators balance chorismate and protocatechuate partition in *Listeria* organisms"

**Supplemental Figures and Data**

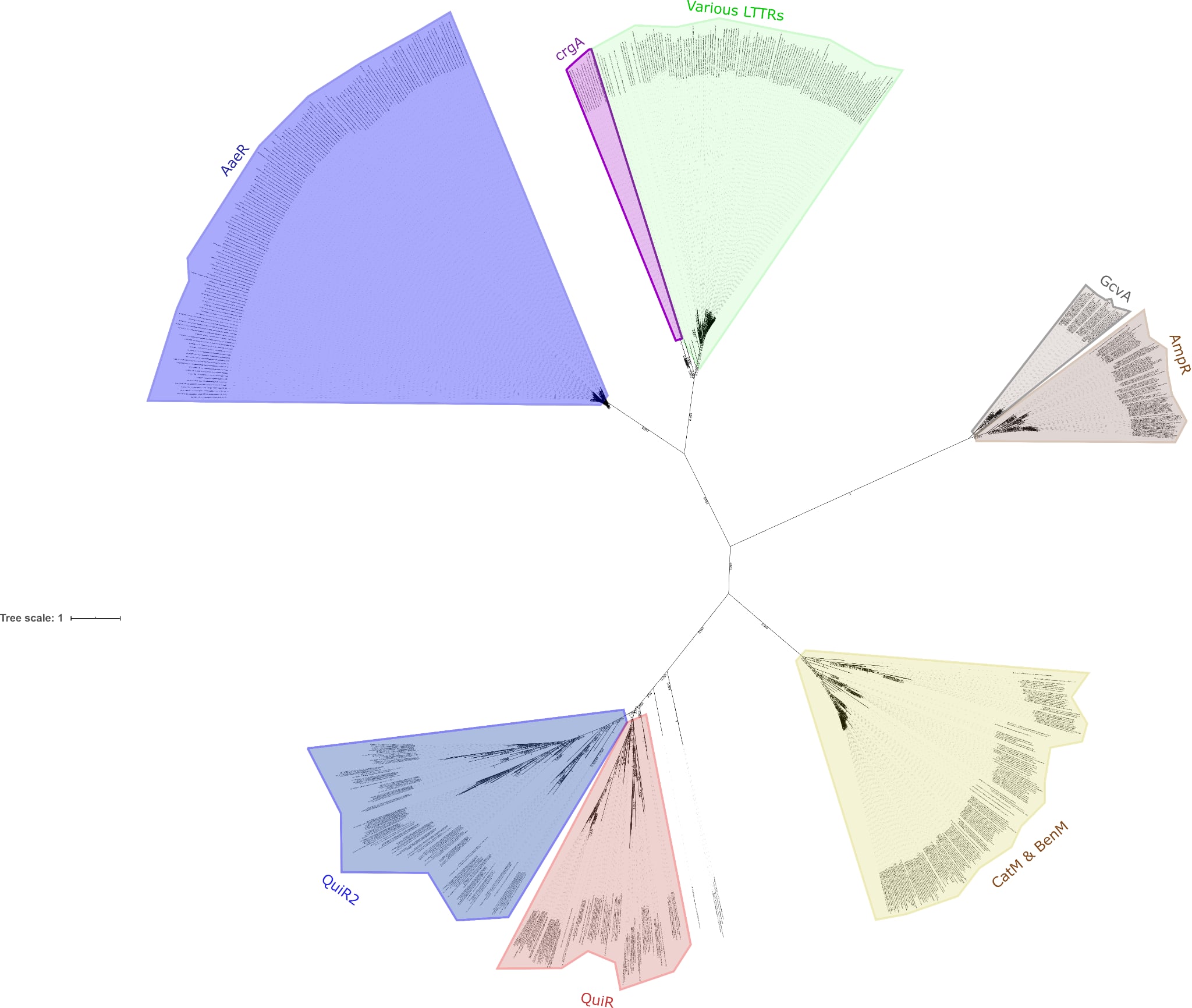

**Supplementary Figure 1.** **Sequences queried from 7 functional LTTR groups: AaeR, CrgA, AmpR, BenM, CatM, QuiR, and QuiR2.** Phylogenetic analyses were performed using the program FastMPTree from XSEDE with 1000 bootstraps. A minimum of 0.81 bootstrap values supports each major branch of the tree. Labeling was performed using ITOL and Inkscape.

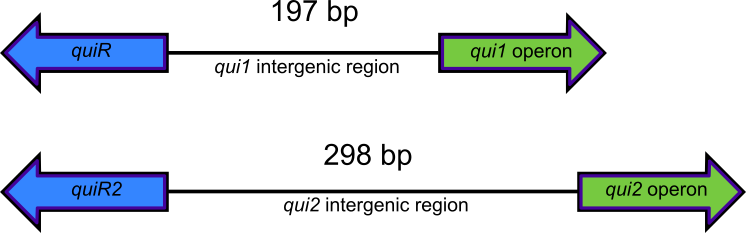

**Supplementary Figure 2. Cartoon schematic showing the intergenic region length of *qui1* (top) and *qui2* (bottom) operons.** The intergenic region is represented by the black line, while the regulators are shown in blue arrow, and the direction of the operons are depicted by the green arrow. The *qui1* operon intergenic region is 197bp in length while the *qui2* intergenic region is 298bp in length.

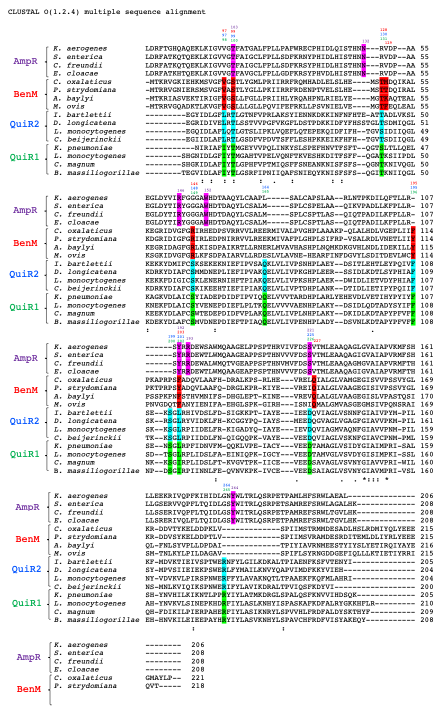

**Supplementary Figure 3: Full amino acid sequence alignments of QuiR2 against 3 distinct families of structurally characterized LTTR regulators**. Represented sequences include the effector binding domain of AmpR, BenM, QuiR2, and QuiR1. Sequences were aligned using the ClustalW Omega software with character counts. Columns with the “.” symbol at the bottom indicates residues that exhibit weak similarity, while the “:” and “*” symbols indicate highly similar and invariant. The active site residues have been highlighted for each corresponding LTTR. Above each colour indicates the position of the residue in the protein.

 
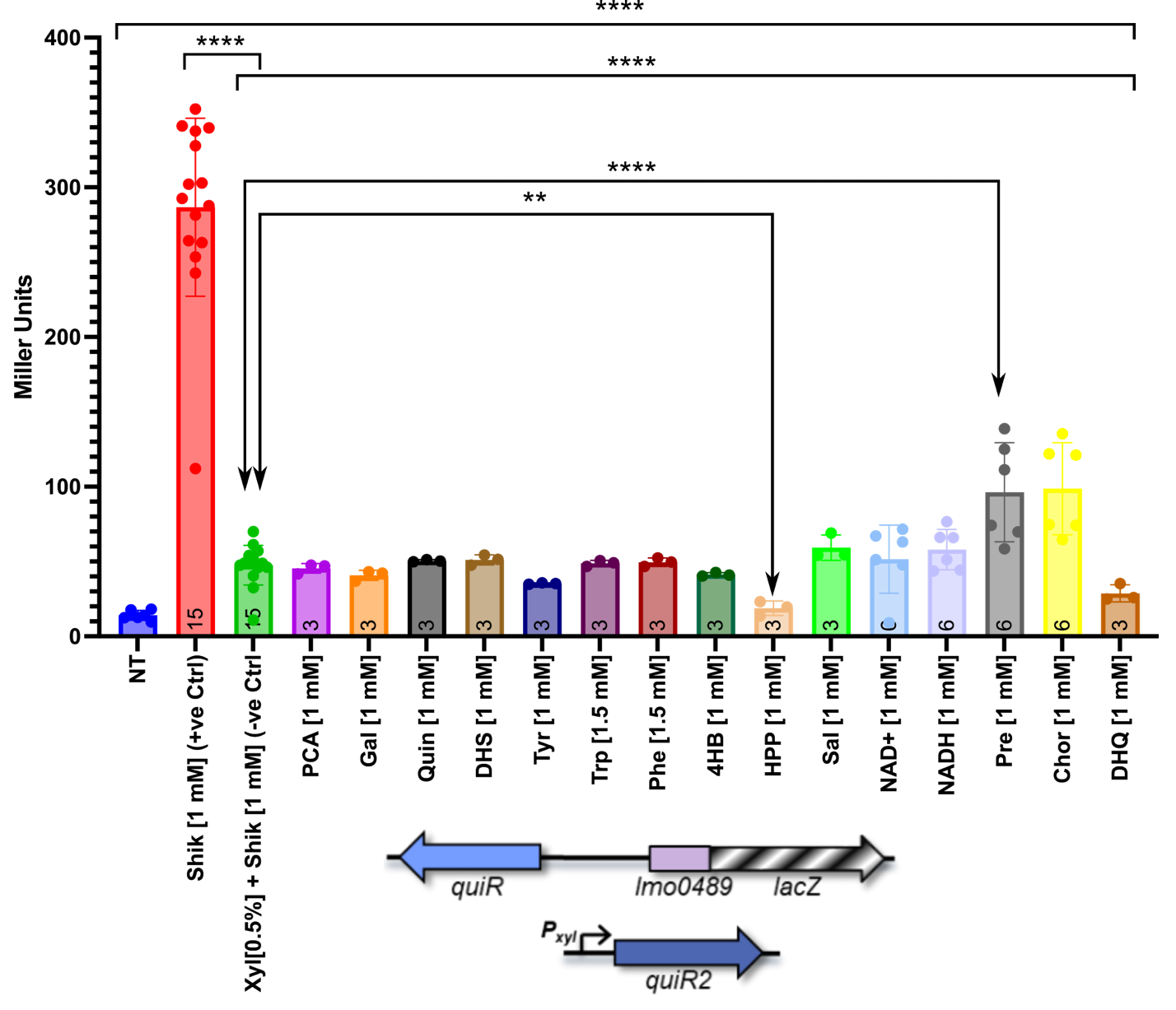

**Supplementary Figure 4. β -galactosidase Assay (Compound Screen): Constructs are visually described with graphics underneath each data bar.** For naming convention, *B. subtilis* transformed with pAX01:QuiR2 (X2), pDG1661:QuiR+qui1 intergenic region (G1IR). All assays were conducted with delayed induction. Replicate counts reflected by data points. NT = no treatment, Shik= shikimate, Xyl= xylose, PCA= protocatechuate, Gal = gallate, Quin= quinate, DHS= dehydroshikimate, Tyr = tyrosine, Trp = tryptophan, Phe =phenylalanine, 4HB = 4-hydroxybenzoate, HPP= hydroxyphenylpyruvate, Sal =salicylate, NAD+= reduced nicotinamide adenine dinucleotide, Pre = prephenate, Chor= chorismate, DHQ= dehydroquinate.

| Species name | Accession # | Sequence region |
| --- | --- | --- |
| *K. aerogenes* | WP_087859132.1 | 84-289 |
| *S. enterica* | WP_061382722.1 | 84-291 |
| *C. freundii* | WP_003839466.1 | 84-291 |
| *E. cloacae* | WP_063941664.1 | 84-291 |
| *C. oxalaticus* | WP_063237961.1 | 81-301 |
| *P. strydomiana* | WP_131319478.1 | 81-298 |
| *A. baylyi* | 2F6G_A | 1-224 |
| *M. ovis* | WP_063514716.1 | 81-299 |
| *I. bartlettii* | WP_022072500.1 | 89-293 |
| *D. longicatena* | WP_028087600.1 | 89-292 |
| *L. monocytogenes* | ECW8281981.1 | 75-278 |
| *C. beijerinckii* | WP_017211209.1 | 89-288 |
| *K. pneumoniae* | VTM40707.1 | 11-215 |
| *L. monocytogenes* | 5TED_A | 1-210 |
| *C. magnum* | WP_066626248.1 | 89-297 |
| *B. massiliogorillae* | WP_042348109.1 | 89-296 |

**Supplementary Table 1. Sequence accessions used for multiple sequence alignment for Figure 4.** The table indicates the species name, the corresponding accession number and the region of the sequence used for the alignment containing the EBD.

| **Compound** | **Binding Energy (kcal/mol)** | **Zinc ID** |
| --- | --- | --- |
| Pinene oxide | -7.4 | ZINC000100019099 |
| (-)-2-Methyl Isoborneol-d3 | -6.6 | ZINC000100302422 |
| Pyrogallol | -5.4 | ZINC000000330141 |
| Isobornyl Formate | -5.4 | ZINC000100254112 |
| 2,3-Dihydroxy-2,4-Cyclopentadien-1-One | -5.4 | ZINC000100828121 |
| Catechol | -5.3 | ZINC000013512214 |
| M-Cresol | -5.1 | ZINC000000897141 |
| 2-Methylcyclohexan-1-one | -5.1 | ZINC000003860605 |
| 2-Methylbutyric Acid cis-3-Hexen-1-yl Ester | -5.1 | ZINC000014438709 |
| Phenolate | -5.0 | ZINC000005133329 |
| 3-Hydroxypyrone | -5.0 | ZINC000001433173 |
| Benzoquinone | -5.0 | ZINC000000895247 |
| Cyclohexanone | -5.0 | ZINC000004528575 |
| Toluol | -5.0 | ZINC000000967534 |
| 2-Aminophenol | -5.0 | ZINC000000157526 |
| Protocatechuate | -4.3 | ZINC000000013246 |

**Supplementary Table 2:** **List of top representative candidate ligands for QuiR2 as identified by Autodock analysis.** The compound name, binding energy (kcal/mol), and the corresponding ZINC IDs are shown.

| **Strains** |  |  |  |
| --- | --- | --- | --- |
| *E. coli* DH5α:pET28MOD: *L.m.quiR2* | *Listeria monocytogenes* EGD-e QuiR2. Kan^r^ | Used to propagate pET28MOD with *Listeria monocytogenes*QuiR2. | Prezioso *et al.* 2018 |
| *E. coli* BL21:pET28MOD: *L.m.quiR2* | *Listeria monocytogenes* EGD-e QuiR2. Kan^r^, Cam^r^ | Used to express *Listeria monocytogenes*QuiR2. | Prezioso *et al.* 2018 |
| *E. coli* DH5α:pET28MOD: *L.m.quiR2 (*Δ89) | *Listeria monocytogenes* EGD-e Δ89 QuiR2. Kan^r^ | Used to propagate pET28MOD with *Listeria monocytogenes*Δ89 QuiR2. | This study^d^ |
| *E. coli* BL21:pET28MOD: *L.m.quiR2 (*Δ89) | *Listeria monocytogenes* EGD-e Δ89 QuiR2. Kan^r^, Cam^r^ | Used to express *Listeria monocytogenes*Δ89 QuiR2. | This study^d^ |
| Bs2 | *Bacillus subtilis*168 Δ *amyE* : [ *lmo0489* (-208 to +38)- *lacZ* ] Cam^r^, Δ *lacA* : empty pAX01 Em^r^ | LacZ fusion strain | This study |
| Bs:XEG2IR | *Bacillus subtilis* 168 Δ *amyE*: [ *lmo2234* (-1181 to 0)- *lacZ*] Cam^r^, Δ *lacA* : empty pAX01:Em^r^ | LacZ fusion strain | This study |
| Bs:X1G2IR | *Bacillus subtilis*168 Δ *amyE* : [*lmo2234* (-1181 to 0)- *lacZ* ] Cam^r^, Δ *lacA* : pAX01:*quiR:* Em^r^ | LacZ fusion strain | This study |
| Bs:X2G1Z | *Bacillus subtilis* Δ *amyE* : [*lmo0489* (-208 to +38)- *lacZ*] Cam^r^, Δ *lacA* : pAX01:*quiR2:* Em^r^ | LacZ fusion strain | This study |
| Bs:X2G2Z | *Bacillus subtilis* Δ *amyE* : [*lmo2234* (-346 to +56)- *lacZ*] Cam^r^, Δ *lacA* : pAX01:*quiR2:* Em^r^ | LacZ fusion strain | This study |
| Bs:X2G1IR | *Bacillus subtilis* Δ *amyE* : [ *lmo0489* (-1187 to +38)- *lacZ*] Cam^r^, Δ *lacA* : pAX01:*quiR2:* Em^r^ | LacZ fusion strain | This study |
| Bs:X2G2IR | *Bacillus subtilis* 168 Δ *amyE*: [*lmo2234* (-1181 to 0)- *lacZ*] Cam^r^, Δ *lacA* : pAX01:*quiR2:* Em^r^ | LacZ fusion strain | This study |
| **Plasmids** |  |  |  |
| pET28MOD:*quiR2* | pET28MOD:His6-*quiR2* | Full Length QuiR2 expression | Prezioso *et al.* 2018 |
| pET28MOD:*quiR2 (Δ89)* | pET28MOD His6‐*quiR2(Δ89)* | Δ89 QuiR2 expression | This study^d^ |
| pAX01 | Plasmid containing a xylose-dependent promoter. *lacA*integration site. Amp^r^, Erm^r^ | Xylose-dependent regulator expression *B. subtilis* for LacZ fusion experiments. | Prezioso *et al.* 2018^a^ |
| pDG1661 | Plasmid containing a *lacZ*fused to an MCS. *amyE*  integration site. Amp^r^, Cam^r‑.^ | Plasmid used to express LacZ during LacZ fusion experiments. Integration screened using starch plates. | Prezioso *et al.* 2018^a^ |
| X1 | pAX01: *quiR:* Em^r^ | pAX01 to express QuiR under xylose induction | Prezioso *et al.* 2018^a^ |
| X2 | pAX01:*quiR2:* Em^r^ | pAX01 to express QuiR2 under xylose induction | This study^a^ |
| G1Z | pDG1661: *lmo0489* (-208 to +38)- *lacZ.*  Cam^r^. | pDG1661 with the intergenic region upstream of *qui1* to assay *qui1* operon expression | Prezioso *et al.* 2018^a^ |
| G1IR | pDG1661: *lmo2234* (-1181 to 0)- *lacZ.*  Cam^r^. | pDG1661 with the intergenic region upstream of *qui1* including *quiR*  to assay *qui1* operon expression when QuiR is natively expressed | This study^a^ |
| G2IR | pDG1661: *lmo2234* (-1181 to 0)- *lacZ.*  Cam^r^. | pDG1661 with the intergenic region upstream of *qui2* including *quiR2*  to assay *qui2 o*peron expression when QuiR2 is natively expressed | This study^a^ |
| G2Z | pDG1661: *lmo2234* (-346 to +56)- *lacZ.*  Cam^r^. | pDG1661 with the intergenic region upstream of *qui2* to assay *qui2* operon expression | Prezioso *et al.* 2018^a^ |

| **Primers** | | | | |
| --- | --- | --- | --- | --- |
| quiR2 (△89): F | CAA*GGATCCAT*GGGGCAGAT TGACTTGGCC | BamH1 | *quiR2* | QuiR2 EBD Cloning |
| quiR2: F | CACTTC*ATTAAT*ATGAATTTACACCATTTACGATACTTCG | AseI | *quiR2* | Full length QuiR2 Cloning |
| quiR2: R | CCTC**CTCGAG**TATACGATGTG CTAACATAAATTG | Xho1 |  | QuiR2 Cloning |
| X1: F | GAATTA*GGATCC*ATGAACTTGCGTCAACTTTACTAC | BamHI^d^ | *quiR* | QuiR expression in *B. subtilis* |
| X1: R | GTAGCA*GGTACC*AGAATTTATGGAGAATATCGAGTATTT | KpnI^d^ | *quiR* |  |
| X2: F | *CCGCGGCCGCG*GTTATATACGATGTGCTAACATAAATTG | SacII | *quiR2* | QuiR2 expression in *B. subtilis* |
| X2: R | TGA*GGATCC*CATGAATTTACACCATTTACGATAC | BamHI | *quiR2* |  |
| G1Z: F | AGTATT*GGATCC*ACAGTTAATGGTTCAAAAATACTCGG | BamHI^d^ | *qui1* promoter | *qui1* regulation |
| G1Z: R | GTCATA*GAATTC*CGCAAATTCATGAAACTACCTCC | EcoRI^d^ | *qui1* promoter |  |
| G1IR: F | AGTATT*GGATCC*ACAGTTAATGGTTCAAAAATACTCGG | BamHI^d^ | *quiR-qui1* promoter |  |
| G1IR: R | GCAAGC*GAATTC*AAAGTTGGTGAGTTATACCGATTTAATG | EcoRI^d^ | *quiR-qui1* promoter |  |
| G2IR: F | GACTC*GAATTC*TTTTATATACGATGTGCTAACATAAAT | EcoRI | *quiR2-qui2* promoter | *qui2* regulation |
| G2IR: R | CAT*GGATCC*AAAAATTATCTCCTCTCCATAATAA | BamHI | *quiR2-qui2* promoter |  |
| G2Z: F | ATT*GGATCC*GTGTAAGAGCTAATCGTGATGGG | BamHI^d^ | *qui2* promoter |  |
| G2Z: R | ATA*GAATTC*CCATGTGCGCCAGTGTGAC | EcoRI^d^ | *qui2* promoter |  |
| OR:F | CTCGGATAATCTGTCATAATAAGCCTCC | None^d^ | *qui1*operon promoter a | P*qui1*EMSA probe |
| OR:R-Biotin | **/5Biosg/**CGCAAGTTCATCGGTTAACCTCCTC | 5' Biotin^d^ | *qui1*operon promoter |  |
| OR2:F | /**5Biosg**/CGCAAGTTCATCGGTTAACCTCCTC | None^d^ | *qui2*operon promoter | Biotinylated P*qui2* EMSA probes |
| OR2:R-Biotin | /**5Biosg**/TCGAGCACGGCCAGGTCGG | 5’ Biotin^d^ | *qui2*operon promoter |  |
| OR2-R | AGACTAAATAGCAGAATACTCACCTG | None^d^ | *qui2*operon promoter | unlabeled P*qui2*EMSA probe |
| pAX:F | TAAGTGTTACCCCTATAAGTTAG | None^d^ | *LacA* | pAX01 integration screen |
| pAX:R | CAAGAACGTTGCTCTAGAG | None^d^ | *LacA* |  |
| lacA:F | GTTGCCGTCATCTTTATTATGC | None^d^ | *LacA* |  |
| lacA:R | GCAAGCGTTTTCATTCTATAG | None^d^ | *LacA* |  |

**Supplementary Table 3. List of bacterial strains, plasmids, and primers used in this study.** ^a^*Listeria monocytogenes* EGD-e strain genomic DNA was used as a template for these primers. Restriction cut sites are italicized, genomic sequences are underlined and mutagenized nucleotides are in bold. ^b^*Bacillus subtilis* strain 168 genomic DNA was used as a template for these primers.^c^ *Listeria innocua* Clip11262 genomic DNA was used as a template for these primers. Restriction cut sites are italicized and genomic sequences are underlined.^d^ View Prezioso *et al.* 2018 for original primers.
